# Increased stress in early life epigenetically rewires pathways related to aging and longevity

**DOI:** 10.64898/2026.09.30.755600

**Authors:** Nelly N. Olova, Tianchi Zhou, Winnie Boner, Robert Gillespie, Edward R. Ivimey-Cook, Riccardo E. Marioni, Emma J. A. Cunningham, Tamir Chandra, Tom J. Little, Pat Monaghan, Davide M. Dominoni

**Author notes:** these authors contributed equally (Pat Monaghan, Davide M. Dominoni).

## Abstract

Environmental conditions experienced early in life have profound consequences for vertebrate health, yet the molecular pathways linking developmental stress to aging and reduced lifespan remain poorly understood. Through a novel highly accurate epigenetic clock, developed from RRBS-based blood DNA methylation profiles, we show that experimental lifespan-reducing early-life stress rapidly accelerates epigenetic age in zebra finch chicks (*Taeniopygia guttata*). Corticosterone-treated 29-day-old chicks share differentially methylated loci with non-treated aged birds and cluster with 6-month-old birds. The transcriptional repressor *ZBTB16* emerges as an epigenetically top age-correlated gene and is strongest affected by corticosterone. Elevated developmental corticosterone causes rapid epigenetic remodelling of metabolic pathways well-established in aging and longevity. Our results demonstrate epigenetic rewiring of the aging trajectory through conserved ‘stress-survival’ mechanisms immediately following elevated early-life stress exposure.

## Introduction

Aging is increasingly recognised as a plastic process, shaped in part by environmental factors experienced throughout life (*1*, *2*). Adverse conditions encountered early during development can have particularly profound long-lasting consequences for health, physiology and lifespan (*3*– *5*), explained with the “allostatic load”, the cumulative “wear and tear” of adaptation to adversity (*6*). Nutritional limitation, psychosocial adversity and physiological stress during early life are increasingly associated with elevated susceptibility to disease, accelerated functional decline and reduced longevity (*1*, *4*, *7*, *8*). However, the mechanisms linking early-life stress (ELS) to altered aging trajectories in vertebrates remain poorly understood (*9*). Much of the existing evidence derives from observational human studies, where early-life exposures are difficult to disentangle from socioeconomic, behavioural and genetic confounders (*10*). Experimental studies remain comparatively rare and have shown variable and contradictory outcomes (*11*). We therefore still lack a mechanistic understanding of how environmental stressors modulate biological aging during development.

DNA methylation (DNAm) has emerged as a powerful biomarker for tracking variation in chronological and biological age (*12*, *13*). Epigenetic clocks leverage highly predictable age-associated changes in DNAm patterns to accurately predict chronological age (*13*). Deviations between predicted epigenetic age and chronological age, commonly termed epigenetic age acceleration, have repeatedly been associated with mortality risk, chronic disease, environmental adversity and stress exposure in humans and other mammals (*14–16*). While epigenetic clocks have been developed for numerous mammalian species (*17*), comparable tools remain scarce in other vertebrates despite their enormous potential for studying the developmental origins of aging.

We addressed these gaps by utilising the zebra finch (*Taeniopygia guttata*), an extensively studied model organism in evolutionary ecology and a powerful system for investigating the long-term consequences of developmental environments (*14–17*). Birds are long-lived endotherms and much of the zebra finch development occurs in the post-hatching nestling phase. Thus, avian models combine relatively long lifespans with experimentally tractable early development, facilitating investigations of how environmental conditions experienced during ontogeny shape aging across the lifespan (*22*, *23*). Avian nestlings are highly amenable to direct experimental manipulation, whilst prior to weaning experimentation on mammals present numerous technical, handling and ethical challenges (*11*).

Experimental elevation of corticosterone (CORT), the main stress hormone in birds, during the nestling stage, was previously shown to heighten the adult stress response (*24*), and reduce adult lifespan in zebra finches by approximately 30% (*25*, *26*). This provides compelling evidence for the utility of this model to address how developmental stress can alter the aging trajectories. We developed a novel, sequencing-based, highly accurate zebra finch epigenetic clock, selected from more than 2000 models, over a cohort spanning the full zebra finch lifespan. We applied the clock to an experimental group of nestlings exposed to increased CORT to examine the proximal effects of developmental stress on age acceleration. Finally, we dissected differential methylation of target genes and driver biological pathways, leveraging methylome changes at the intersection between CORT treatment and the natural aging trajectory.

## Results

### Improved sequencing-based regional age prediction

Our AGE cohort (clock training set) numbered 99 red blood cell (RBC) samples, spanning over a range of 9 years (3391 days), with a balanced proportion of males and females (Fig. 1A). Additional 96 samples were used for independent testing of the clocks (test set), with a more limited age range. The CORT cohort comprised 96 nestlings including controls (Fig. 1A, Table S1). Mean global DNAm of the training set was 20.34% and no difference was observed between processing LOTs (Fig. S1A, B). As expected for Reduced Representation Bisulfite Sequencing (RRBS) data, calls were exclusively localized in CpG (cytosine-phosphate-guanine) islands (CGIs) and within five kilobases (kb) of gene bodies and promoters (Fig. S1C).

**Figure 1.**
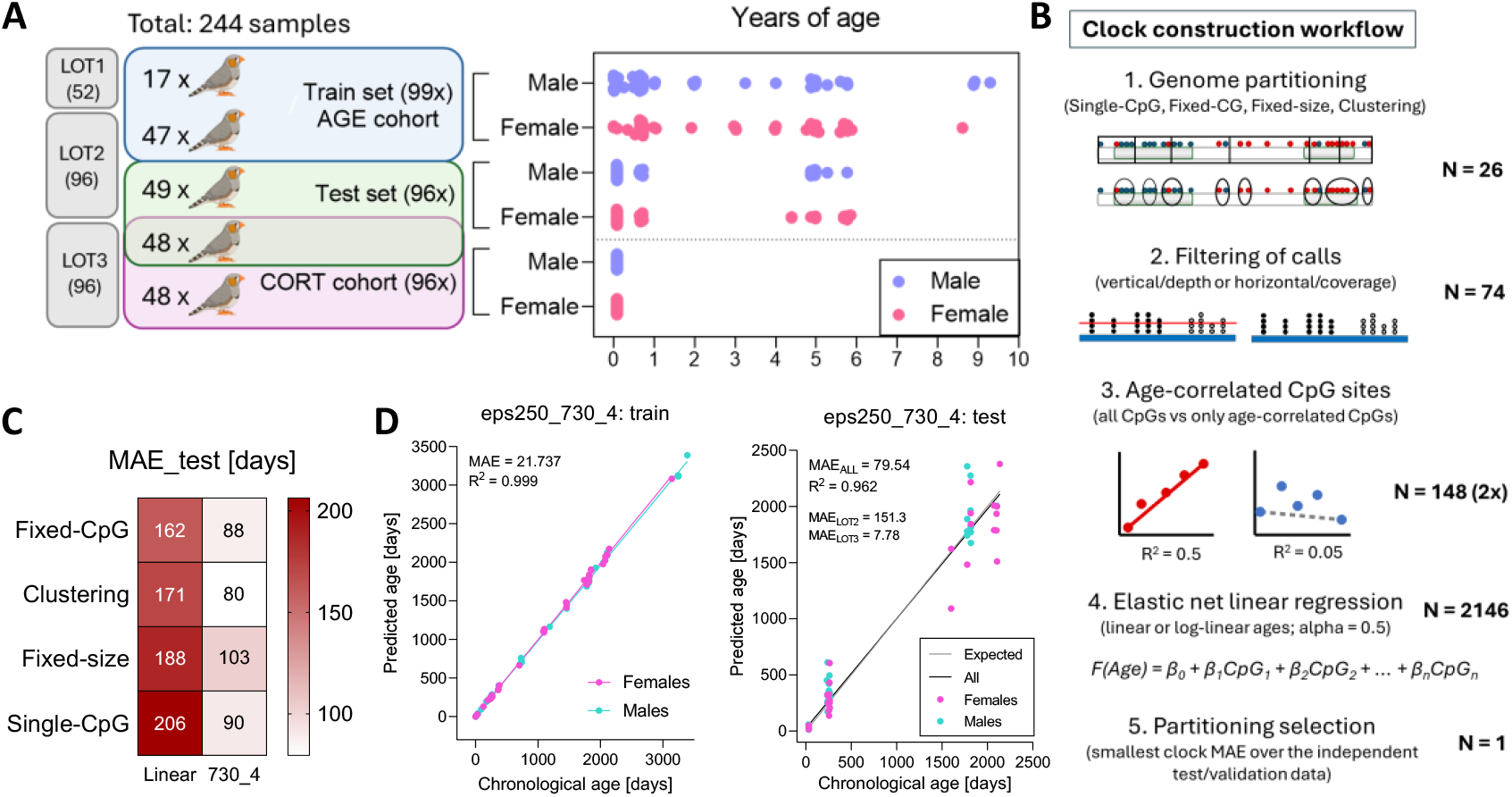
Data structure and RRBS-based clock construction. (**A**) 244 blood (RBC) samples from 209 healthy birds derived from three experimental and sample processing LOTs, as detailed in Methods. None of the birds used for clock building and validation had been subjected to treatments that affected mortality patterns. Right panel represents the age and sex distribution of 244 zebra finch samples within the train set (AGE cohort), test set and CORT cohort. (**B**) The clock construction workflow: 1. Data preparation and genome partitioning; 2. Filtering of calls and CpG positions; 3. Selection and removal of non-age-correlated CpGs; 4. Clock training (n=99) via elastic net linear regression, with and without logarithmic transformation of ages below 730 days (two years), and applying the 2,146 final models to the independent test set (n=96); 5. Final model selection across the 2,146 partitioning and filtering combinations, based on the lowest mean absolute error (MAE) on the test set. (**C**) Top performing linear and log-linear models on the independent test data across the four partitioning approaches. (**D**) Performance of the top clock over the train and test data; eps250 derives from the cluster size of the Clustering approach used and 730_4 points to 730 days of log-linear age cutoff and four days added as a coefficient. MAE of eps250_730_4 on LOT2 birds is 151.3 days and on LOT3 controls is 7.78 days.

We recently demonstrated that averaging methylation over regions, rather than using individual CpG sites, can significantly improve clock accuracy and transferability across RRBS data (*27*). To identify the best region size and parameters, we utilized a comprehensive clock-construction workflow across four different data-partitioning approaches and varied several parameters in parallel (Fig. 1B) (*28*). Alongside the classical individual CpG approach (‘Single-CpG’), we used: sliding windows of a fixed genomic size (‘Fixed-size’), density-based clustering of CpGs (‘Clustering’), and a fixed number of CpGs (‘Fixed-CpG’) (Fig. S2A). We selected the top performing clock based on the lowest mean absolute error (MAE) over the test set (Fig. 1B).

We generated both linear and log-linear clocks, after log-transforming young ages to accommodate for non-linear rates of methylation changes during organismal development, as commonly done for mammals (*14*, *17*) (Fig. S2 B,C). Log-linear clocks outperformed the linear clocks with a two-fold reduction in MAE (Fig. 1C). The top performing clock, selected for further use, was derived from the Clustering approach (‘eps250_730_4’, MAE = 80 days) (Fig. 1D, Table S2). We found little influence of repeated sampling on the predicted epigenetic age and detected no significant technical batch effect (*p* = 0.662).

### The epigenetic clock sites enrich for developmental, circadian and immune processes

The top performing ‘eps250_730_4’ predictor covered mostly CGIs that localised within gene bodies and less in predicted promoters compared to the genomic average (Fig. 2A and Fig. S3A). The majority of eps250_730_4 sequences showed higher methylation with age (62% overall, Fig. 3B and 54% to 65% across features, Fig. S3B, C), contrary to the mouse regional RRBS clock (*27*). Clock segments were distributed across most chromosomes, including microchromosomes and the Z sex chromosome (Fig. 3C and Fig. S3C). Nearly all loci mapped within 5kb of a gene to a total of 102 genes, out of which 84 have known orthologs (Fig. 3C, Table S2).

**Figure 2.**
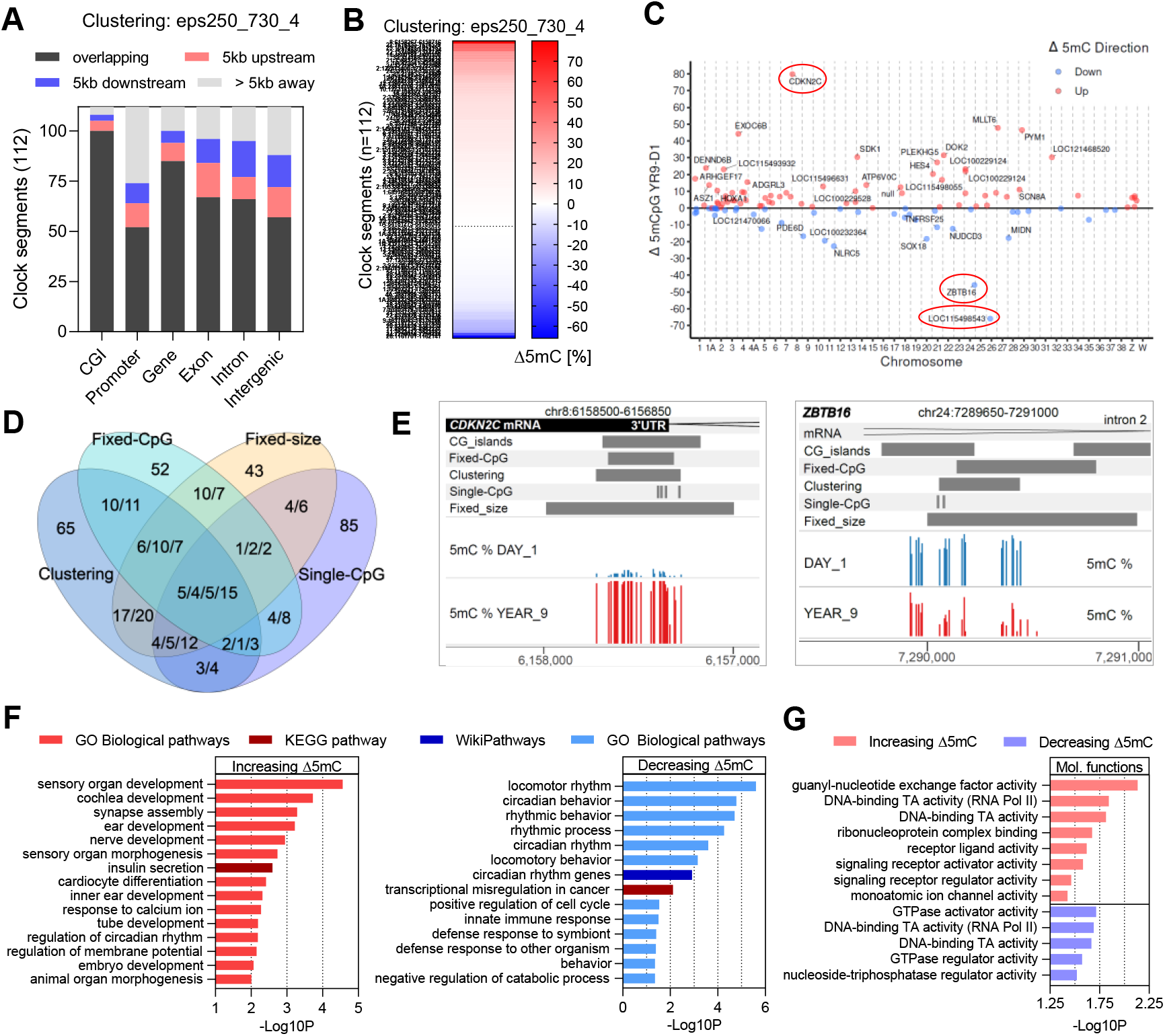
Structural and functional features of the zebra finch epigenetic clocks. (**A)** Genomic feature mapping of the top log-linear Clustering eps250_730_4 clock: each bar represents the proportion of clock segments that overlap, are within 5kb or beyond 5kb from the feature of interest. (**B)** Clock segments of the eps250_730_4 clock with higher and lower methylation change (Δ5mC) with age; the values represent the difference between ‘day 1’ and ‘year 9’ samples. (**C)** Methylation change of all 730_4_eps_250 regions across their chromosomal locations; labels represent annotated genes for all regions with methylation change above 10% in both directions (Δ5mC > |+/-10|). (**D)** Overlapping sequences between the four top log-linear zebra finch clocks per partitioning approach. Each number corresponds to the number of segments from one approach, because a segment from one regional approach can overlap with multiple segments from another approach; order from left to right: Clustering, Fixed-CpG, Fixed-size, Single-CpG. (**E)** Chromosomal view of the top overlapping clock segments over *CDKN2C* (left) and *ZBTB16* (right). Clock segments and CGIs are depicted as grey bars, the mRNA is in black, and methylation percentage (5mC %) bars are in blue for ‘day 1’, and in red for ‘year 9’ samples. (**F)** Pathway enrichment analysis and biological processes gene ontology for the hypo- and hyper-methylated regions from the eps250_730_4 clock. (**G)** Molecular functions enrichment for the hypo- and hyper-methylated regions from the eps250_730_4 clock; TA stands for “transcriptional activator”.

**Figure 3.**
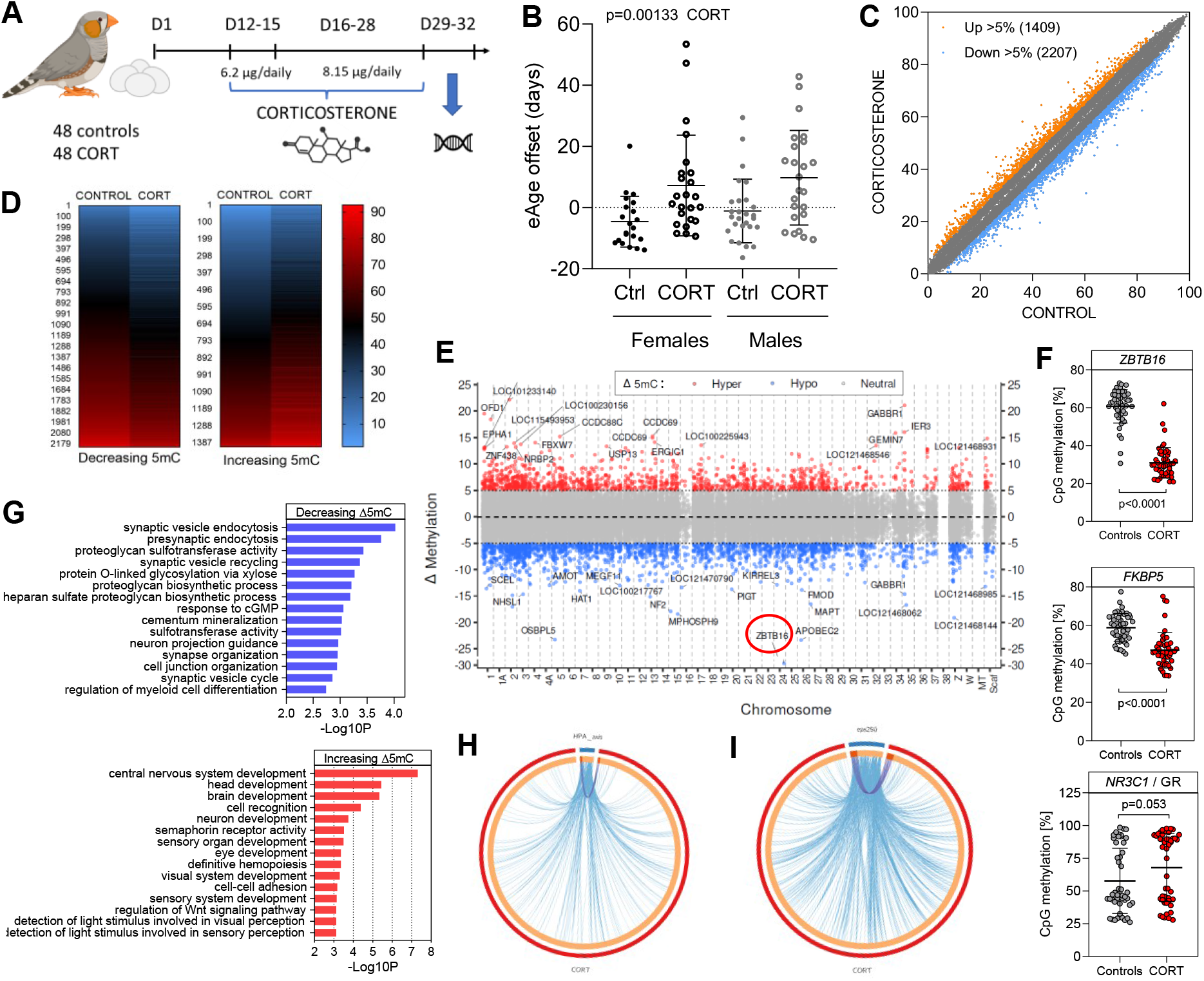
Early life CORT supplementation accelerates epigenetic age and severely demethylates *ZBTB16*. (**A**) The CORT experiment was conducted as follows: starting on day 12 (D12) post-hatching, 48 zebra finches were orally supplemented with CORT twice daily, mimicking natural elevations of CORT due to stress; controls (n=48) were given only the carrier oil (*20*). Treatment lasted for 17 days, up to and including day 28 (D28). Whole blood was collected on day range 29-32 (D29-32). (**B**) Accelerated epigenetic age of 11 days on average in the CORT-supplemented birds, for both sexes (n = 24 each); ‘Ctrl’ stands for ‘controls’ (21 females, 27 males). Significance was assessed with a linear mixed effects model, after correcting for family (nest) (Table S5). Error bars represent standard deviation (S.D.). (**C**) A logistic regression test identified 60,885 differentially methylated regions (DMRs) with p < 0.01 (depicted in grey), out of which only DMRs with an absolute mean methylation change of more than 5% across the CORT group were considered as biologically meaningful: 2207 hypo-methylated (in blue) and 1409 hyper-methylated regions (in orange). (**D**) Mean methylation of the 3616 hyper- and hypo-methylated DMRs across the control and CORT groups. (**E**) Chromosomal distribution of the 1409 hyper- and 2207 hypo-DMRs, with top 20 genes labelled on each side; *ZBTB16* is circled in red. (**F**) Methylation changes of the top CORT-affected gene *ZBTB16* (Δ5mC_mean_=30%), the co-chaperone *FKBP5* (Δ5mC_mean_=11.7%) and the glucocorticoid receptor (GR) *NR3C1* (Δ5mC_mean_=10%, borderline n.s.); error bars represent S.D., plot p-values are derived by a t-test, however, all DMRs showed high significance (p<0.0001) with a logistic regression test. (**G**) Top 15 highly significant GO terms associated with the hyper- or hypo-methylated CORT-affected genes. (**H**) A circos plot depicting direct (shared gene = purple lines) and indirect (shared biological pathway = blue lines) overlap between 1,441 known-function CORT-affected genes and 35 HPA-axis genes; Fisher’s p=0.021. (**I**) A circos plot depicting the direct and indirect functional overlap between 1,441 CORT-affected genes and the 84 known-function eps250_730_4 clock-associated genes; Fisher’s p=0.00647, FDR=0.0129.

To identify top gene hits, we looked at coordinate overlaps between all four log-linear approaches (Fig. 1D). Five genes mapped to overlapping regions from all four top clocks (Fig. 3D). The top hit, covered by 9 CpGs from the Single-CpG clock, was an unknown-function non-coding (nc)RNA, *LOC115498543* (15kb away from *CR2*), followed by *CDKN2C*, *ZBTB16*, *BCL2L11* and *ADAP2* (Fig. 3E and Fig. S3D). Gene ontology (GO) of eps250_730_4 clock-associated genes showed enrichment for organ morphogenesis and development terms in the hyper-methylated group. The hypo-methylated genes showed a strong signature of rhythmic processes and immune response (Fig. 3E, Table S3). Molecular functions were dominated by signalling and transcription factor activities in both directions (Fig. 3G).

### Early life CORT exposure induces accelerated age and strongly demethylates ZBTB16

We next explored if we could capture accelerated epigenetic age in the nestlings subjected to early-life CORT treatment (Fig. 3A), known to substantially reduce life expectancy (*20*). We observed net acceleration of 11 days for the CORT group, irrespective of sex (*p* = 0.0013, Fig. 3B and Tables S4, S5), which persisted four days following treatment (Fig. S4A). Global genomic methylation did not display any differences and showed good correlation between treated and control birds (Fig. S4B-D).

We identified 3,616 differentially methylated regions (DMRs, p<0.01, FDR<0.03) with absolute mean methylation changes between 5% and 30% for the treated group (Fig. 3C-E and Fig. S4E). Genomic feature distribution followed the typical RRBS pattern (Fig. S1C), but was less enriched in predicted promoters and CGIs, and more within intergenic sequences (Fig. S4F). The CORT-DMRs mapped to 1,879 genes, out of which 1,441 had known human orthologs (Fig. 3E and Table S6). The most significantly differentially methylated gene was *ZBTB16* (Δ5mC = - 30%, Fig. 3E, F), followed by *APOBEC2, OSBPL5, GABBR1* and more (Fig. 4E and Fig. S5A,B). The FK506-binding protein 5, encoded by *FKBP5* and previously implicated in stress response dysregulation (*29*), was also significantly demethylated (Δ5mC = −11.7%). The glucocorticoid receptor (GR) *NR3C1* was among the DMRs, albeit highly variable across individuals (Δ5mC = 10%, Fig. 4F). Whilst *FKBP5* showed recovery on days 3 and 4 post CORT exposure, *ZBTB16* maintained robust hypo-methylation (Fig. S5C). Gene ontology (GO) analysis showed a marked enrichment in nervous system and sensory development processes within the hyper-methylated genes, and in synaptic plasticity and extracellular matrix remodelling processes within the hypo-methylated genes (Fig. 4G and Fig. S6). А direct comparison with 35 hypothalamic-pituitary-adrenal (HPA) axis-associated genes (*30*, *31*) revealed six shared genes, which wasn’t a significant enrichment (p=0.21), but showed a wide functional overlap (Fig. 4H, Fig. S4G, Table S3). However, a highly significant direct gene overlap (18, p = 0.0065, FDR=0.013) and a much wider overlap of biological pathways was observed with the eps250_730_4 clock genes, selected for their log-linear age-association and predictive power to track age (Fig. 4I, Fig. S4G).

**Figure 4.**
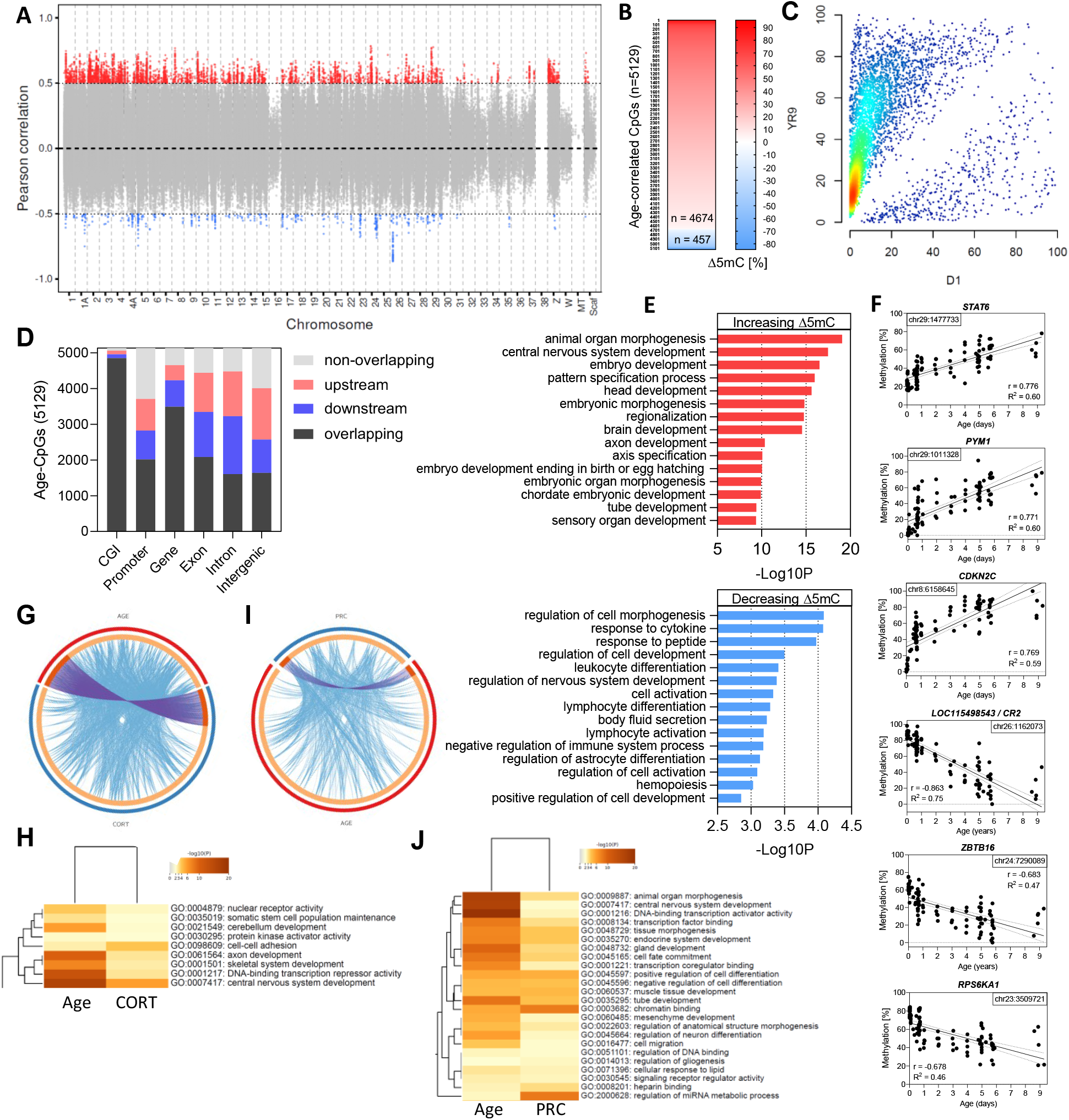
Genome-wide age-associated CpGs functionally overlap with CORT-loci. (**A**) Age-CpGs were selected by applying a Pearson correlation coefficient cutoff r > |+/-0.5| to all shared CpGs in our AGE-cohort. (**B**) Direction of age-associated methylation change (Δ5mC) of all 5,129 Age-CpGs. (**C**) Mean methylation values of all 5,129 Age-CpGs between 1-day-old (D1) and the oldest 9-year-old birds (9YR). (**D**) Genomic feature enrichment of the 5,129 Age-CpGs. (**E**) GO enrichment analysis of higher (4,671) and lower (458) in methylation Age-CpGs. (**F**) Methylation values of selected top three hyper- and hypo-methylated age-correlated loci across the AGE-cohort; only the top-correlated CpG per each gene is shown, out of 47 Age-CpGs mapping to *PYM1*, 6 to *STAT6*, 34 to *CDKN2C*, 44 to *LOC115498543*, 40 to *ZBTB16* and 12 to *RPS6KA1*. (**G**) A circos representation showing directly overlapping (purple) and functionally overlapping (blue) lines for 851 known-function genes mapping to the Age-CpGs (AGE) and the 1,441 known-function CORT-affected genes (CORT); Fisher’s p=3×10^-12^, FDR=1.2×10^-11^. (**H**) Statistically significant GO terms of shared pathways between the age-associated and CORT-affected genes. (**I**) Directly overlapping (purple) and functionally overlapping (blue) lines for 851 genes mapping to the Age-CpGs (AGE) and 339 PRC-target genes (PRC); Fisher’s p=0.222. (**J**) Statistically significant GO terms of shared pathways between the age-associated and PRC-target genes.

### Age-associated CpGs show positional and functional overlap with CORT-affected loci

To further decipher the connection between CORT-affected genes and the lifespan-reducing phenotype, independently of the epigenetic clock results, we investigated genome-wide age-correlated CpGs and their features. Genome-wide average CpG methylation (Fig. S1B) showed weak but significant positive correlation with age (r=0.39, R^2^=0.16, p<0.0001) (Fig. S7A). We calculated Pearson correlation for all shared CpG sites for the AGE-cohort and selected 5,129 ‘Age-CpGs’ with |r| ≥ 0.5 (Fig. 4A, Fig. S7B, Table S7), which were distributed across most chromosomes, with the exception of chr38 (Fig. 5A). The vast majority (91%) showed an increase in methylation with age (Fig. 4B,C) and were exclusively located within CGIs (Fig. 4D, Fig. S7C,D). Pathway enrichment of 851 Age-CpG-mapped genes was dominated by developmental categories, where organismal and nervous system development and morphogenesis characterised the hyper-methylated genes, whilst cellular differentiation, morphogenesis and activation, with a pronounced focus on blood cells, dominated the hypo-methylated genes (Fig. 4E, Fig. S8). The top age-correlated loci mapped to *STAT6*, *PYM1* and *CDKN2C* (increasing), and the unknown-function lncRNA *LOC115498543* (*CR2*), *ZBTB16* and *RPS6KA1* (decreasing in methylation). The positional and functional overlap with CORT-affected genes revealed a highly significant 165 shared genes between the two groups (p<0.0001, FDR<0.0001), and the rest were highly associated with the same biological pathways (Fig. 5G, H).

**Figure 5.**
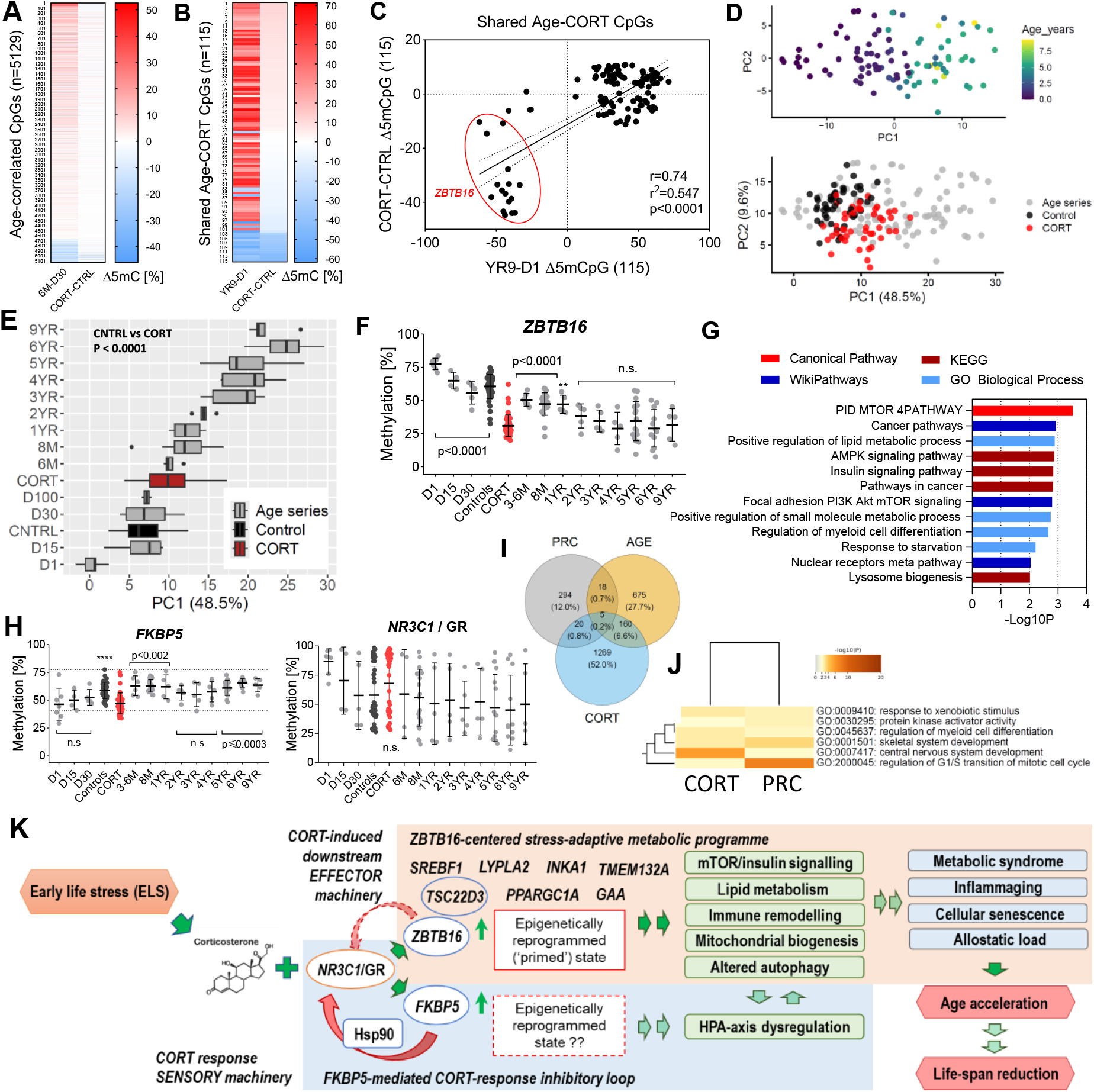
CORT remodels epigenetically *ZBTB16* and rewires core longevity-associated pathways. (**A**) Methylation changes in 5,129 Age-correlated CpGs between 6-month-old (6M) and 30-day old birds (D30) alongside their changes induced by early-life CORT exposure (CORT-CTRL). (**B**) Methylation changes in 115 shared Age-CORT CpGs for the entire 9-year life-span (YR9-D1) and induced by early-life CORT exposure (CORT-CTRL). (**C**) Correlation of methylation changes (Δ5mC) of the 115 shared Age-CORT CpGs; CpGs belonging to *ZBTB16* are circled in red. (**D**) PCA of the 115 shared Age-CORT CpGs: the age series trajectory was defined by our AGE-cohort (99 clock-training samples, upper panel), and the CORT-cohort samples were projected onto that PC space (lower panel). (**E**) PC1 mean values per age group, with SD, presented for the age categories in the AGE-cohort (age-series) and the experimental CORT and Control groups. (**F**) Methylation of the most highly affected 370-bp long *ZBTB16* CORT-DMR (Chr24:7290035-7290405) across the age trajectory (grey), and upon early life CORT exposure (CORT=red, Controls=black); error bars represent SD. (**G**) GO terms enrichment of the 26 genes annotated to the 115 Age-CORT CpGs, derived from different ontology databases. (**H**) Methylation of the *FKBP5* and *NR3C1*/GR, across the age trajectory (grey), and upon early life CORT exposure (CORT=red, Controls=black); error bars represent SD. (**I**) Overlap between age-associated genes, CORT-affected and PRC-target genes; gene numbers and their percentage are labelled at each overlap. (**J**) GO enrichment terms for 29 shared genes between early life CORT exposure and PRC targets. (**K**) A proposed model for two parallel programs (stress-adaptive metabolic and CORT-response inhibitory) derived from the epigenetically affected genes and remodelled biological pathways immediately following a prolonged early-life CORT exposure. Dark green arrows point to results shown in this study, light green and red arrows point to outcomes deducted from the literature.

To further explore functional conservation with mammals, we investigated associations with known Polycomb repressive complex (PRC) target genes (*32*) (Table S3). We found 27 shared genes, which wasn’t significant (p=0.222) but showed strong functional GO term overlap (Fig. 5I,J).

### CORT acts on key longevity pathways to rewire epigenetic aging

To further explore the relationship between Age-CpGs and CORT exposure, we asked if Age-CpGs change methylation in the same direction in CORT individuals as they do in aged individuals. Despite no pronounced shifts in overall 5mC distribution (Fig. S9A), 2,186 sites (47%) out of 4,671 hyper-methylated Age-CpGs increased their methylation, while 301 (66%) out of 458 hypo-methylated Age-CpGs decreased their methylation with the CORT treatment (Fig. 5A and Fig. S9B). There was a weak but significant positive correlation between the two groups (Pearson r=0.19, p<0.0001, Fig. S9C).

To narrow down the effect, we looked at the intersection between Age-CpGs and CORT-DMRs and found 115 shared Age-CORT CpGs (Fig. 5B, Fig. S9D, Table S8). Pearson correlation of Δ5mC of the Age-CORT CpGs was very high (r = 0.74) and gene annotation revealed that an entire highly hypo-methylated group of 16 CpGs mapped to *ZBTB16* (Fig. 5C). Other genes with multiple CpG changes mirroring natural aging were *PLEKHG5*, *ZNF532*, *ARSA*, *ALKBH4* (higher Δ5mC) and *GSTT2B* (lower Δ5mC) (Fig. S10A). We then mapped the Age-CORT 5mCpG changes with a principal component analysis (PCA), where PC1 captured the aging trajectory (r_age(PC1)_=0.81, r_age(PC2)_=-0.04), accounting for 48.5% of the variance (Fig. 5D, Fig. S10B,C). Projecting the 29-day-old CORT-cohort methylation values onto the aging trajectory PC space, placed the CORT samples shifted to the right from Controls (p<0.0001, Fig. 5D) and next to the 6-month-old birds, while the Controls placed next to the 30-day-old group (Fig. 5E). The CORT group showed high variation among individuals with some sitting with 5-year-old birds. Indeed, *ZBTB16*’s CORT-induced hypo-methylated levels were indistinguishable from aged (above two-year-old) birds’ and significantly differed from younger (one-year-old and less) birds’ levels (p<0.0001, Fig. 5F). GO enrichment of processes and molecular functions of the associated 26 cross-sectional Age-CORT genes highlighted key nutrient metabolism and autophagy-related pathways, such as mTOR/AMPK and insulin signalling, lipid and lysosomal biogenesis, which have been long established in longevity research as modulators of aging and lifespan (*25*) (Fig. 5G). Key genes associated with these metabolic pathways showed hypo-methylation and include *SREBF1*, *PPARGC1A*, *TSC22D3*, *IKA1*, *LYPLA2* and *GAA* (Fig. S10A). Unlike *ZBTB16*, *FKBP5*’s CORT-induced hypo-methylation (Fig. 3F) did not follow its aging trajectory, and comparably low levels were only observed at day 1 (Fig. 5H). Both the GR, *NR3C1*, and the mineralocorticoid receptor (MR), *NR3C2*, showed lower average hyper-methylation (Fig. 3F, Fig. S10D). However, those values were not significant in the context of their variable levels throughout the natural aging trajectory (Fig. 5H, Fig. S10D). Finally, we looked at the possibility of PRC regulation on the CORT-responsive genes. There was no significant shared target gene enrichment (p=0.97), but the 29 shared genes showed functional coherence in the myeloid, skeletal and nervous system development (Fig. 5I, Fig. S10E). Five of those were shared with the natural aging gene list (p= 0.0438) (Fig. 5J).

## Discussion

Experimental elevation of glucocorticoid exposure during the nestling stage, as would occur in stressful environmental circumstances, permanently elevates zebra finch stress responsiveness, at the price of shortened adult lifespan (*20*). By examining changes in DNA methylation patterns, we reveal that 17 days of CORT treatment significantly accelerated epigenetic age, prior to any measurable change in mortality risk. We further identified a CORT-responsive methylome signature, which adopts an aged-like state upon early life stress (ELS), positioning 29-day-old chicks at around six months of age. Our findings suggest that adverse developmental conditions rapidly embed in the blood methylome shifting it toward an aging-associated state. We further reveal candidate molecular pathways across the nutrient and energy metabolism spectrum, such as mTOR/AMPK, lipid regulation and insulin signalling, which link the conserved stress-survival response of ELS with the accelerated biological aging and reduced lifespan phenotypes (*20*), supporting recent evolutionary theories on why organisms age (*3*, *13*).

We developed and validated a novel epigenetic clock for the zebra finch, an established model organism for studying the long-term impacts of ELS (*18*, *24*). We used RRBS-derived DNA methylation profiles from blood (RBC), as a molecular readout of a systemic rather than tissue-specific response (*9*). We employed a comprehensive workflow, evaluating more than 2000 data preparation and analytical strategies for RRBS-based clock construction (*28*), which provided us with the necessary sensitivity to apply the predictor on 29-day-old chicks immediately after experimentally induced ELS. Our observation establishes that epigenetic clocks can successfully capture accelerated epigenetic age for avian species similarly to mammals.

Our differential methylation analyses extended to the intersection of CORT-treatment and normal aging, uncovering a shared methylation signature of 115 CpGs and associated candidate molecular pathways. Shared target CpGs changed methylation in the same direction across ELS and age, demonstrating an immediately aged methylome at key loci, independently of age predictors.

*ZBTB16* is the core element of the shared Age-CORT methylation signature: it is the single most demethylated gene, appearing unequivocally in top clock targets, CORT-DMRs and age-correlated CpGs. *ZBTB16*’s overexpression is a documented highly conserved stress response across mammals, birds and fish (*34*, *35*). However, it has never been directly implicated in the pathophysiology of ELS (*35*, *36*). Conversely, the GR/*NR3C1* and its negative regulator and co-chaperone *FKBP5,* are glucocorticoid-response genes known to be differentially methylated after ELS and implicated in the adverse physiological effects in child- and adulthood (*35*, *36*). Our results confirm differential methylation of *FKBP5* and a non-significant elevation for *NR3C1*/GR immediately following ELS.

*ZBTB16* was named the ‘glucocorticoid memory’ gene, shown to sustain a permanent transcriptionally primed CORT-responsive state after initial stimulation (*35*, *37*). We show that, following repeated CORT elevation, blood *ZBTB16* in 29-day-old chicks has rapidly reached its most demethylated ‘aged’ levels within the life-course trajectory. Importantly, prolonged CORT stimulation is known to cause stably reprogrammed epigenetic states of target genes, which fail to recover post stimulation (*37*). While *FKBP5*’s 5mC levels start recovering after day 3 post CORT withdrawal, this is not evident for *ZBTB16*. Our data therefore suggest that stable demethylation of *ZBTB16* could be responsible for its ‘stress-memory-gene’ transcriptionally primed state (*35*, *37*).

*ZBTB16* is a direct CORT target via the GR/*NR3C1*, and a potent transcription factor with wide-reaching downstream effector functions, including in cellular differentiation and energy metabolism, immune response, autophagy, haematopoiesis and cancer (*35*, *37*). It has previously been linked to key molecular hallmarks of metabolic syndrome, including direct control of adipogenesis, mitochondrial biogenesis, lipid oxidation, carbohydrate metabolism and insulin sensitivity (*38*). In the context of our intersected Age-CORT signature’s pathway enrichment results, this points to a model, where the immediate effect of ELS is a rapidly activated metabolic and immune stress-survival programme, centred around the epigenetic remodelling to an aged-like state of *ZBTB16*. Post-ELS birds have increased stress responsiveness (*20*), suggesting that the survival metabolic programme could be stably ‘primed’ by the ‘stress-memory-gene’ *ZBTB16* to contribute to an increased allostatic load accumulation, ultimately resulting in an accelerated aging trajectory and substantially reduced life expectancy (Fig. 5K). Altered metabolic phenotypes are commonly displayed post-ELS in humans and rodents, such as obesity, metabolic syndrome, or type 2 diabetes (*39*, *40*).

Our data do not convincingly support a rapid repression of the GR/*NR3C1* in RBC following ELS, neither any plausible role for its hypermethylation, given its highly variable methylation profile across non-treated individuals of all ages. In this scenario, the conserved phenotype of reduced CORT sensitivity via hypermethylation of *NR3C1* could be the outcome of a compensatory feedback mechanism in response to the activated and ‘primed’ by *ZBTB16* metabolic programme, to limit the extent of allostatic load accumulation (Fig. 5K). A hypothetical role for *ZBTB16*, being a potent transcriptional repressor, capable of inducing lasting transcriptional repression of its target genes, cannot be excluded in this (*41*). Thus, our results represent a conceptual shift from the current paradigm, where a reduced CORT sensitivity and lasting HPA axis dysregulation through hypermethylation of the GR/*NR3C1*, are recognised as the major driver, rather than a compensatory consequence, of the post-ELS metabolic programme and stress-induced allostatic load (*42*).

A strong conservation is observed in the post-ELS pathophysiology between birds (*29–31*), mammals (*45*, *46*) and other vertebrates (*4*), pointing to a likely evolutionary conservation of the observed epigenetic remodelling of the stress-response and survival mechanism. Our results point to a coherent biological mechanism with gene players and pathways, previously independently implicated in ELS or aging across vertebrates (*5*, *26*, *45*). Furthermore, the genomic composition of the zebra finch clocks revealed features broadly consistent with findings from mammalian epigenetic clocks, whilst also reflecting distinctive aspects of avian genome organisation. Bird genomes are compact and gene-rich, containing short intergenic regions and fewer repetitive elements (*26*, *27*), which explains the strong enrichment of clock regions and Age-CpGs within genes and promoters, relative to mammals, where repetitive elements often constitute major targets of age-related methylation drift (*27*).

Age-dependent hypermethylation at CpG islands, usually associated with PRC-regulated regions, is among the most reproducible signatures of vertebrate aging, linked to altered transcriptional regulation, stem cell function and developmental gene control (*9*, *28*). The functional association with PRC targets we observed, suggests that core aspects of development and epigenetic aging are evolutionarily conserved across vertebrates.

Our observations show how adverse early-life environments induce long-lasting physiological changes that influence later-life health and lifespan (*49*). Leveraging birds as a particularly powerful system for testing hypotheses about the developmental origins of aging, our work offers a novel perspective on our mechanistic understanding of how ELS response develops to impact adult health, aging and lifespan.

## Supporting information

Supplementary Table S3 - Gene lists

Supplementary Table S6 - Annotated coordinates of CORT-DMRs

Supplementary Table S7 - Annotated coordinates of Age-CpGs

Supplementary Table S8 - Shared Age-CORT CpGs

Supplementary Table S1 - Sample methylation and metadata

Supplementary Table S2 - Annotated coordinates and weights eps_250_730_4

## Acknowledgments

The authors thank the Edinburgh Clinical Research Facility’s Genetics Core staff for excellent sequencing services.

## Funding

Natural Environment Research Council grant NE/S005773/1 (DMD)

Reinvigorating Research Grant of the University of Glasgow (DMD)

European Research Council Advanced Grants No. 268926 and 101020037 (PM)

BBSRC Fellowship UKRI899 (NNO)

BBSRC Research Grant BB/P009174/1 (PM)

University of Edinburgh School of Biological Sciences Seed Funding Award (NNO)

MRC Doctoral Training Programme in Precision Medicine (TZ)

UoE Chancellor’s Fellowship (TC)

## Author contributions

Conceptualization: PM, NNO, DMD

Data collection: NNO, WB, TZ, PM

Data curation: NNO, DMD, PM

Methodology: NNO, TZ, WB, RG, ERIC, TJL, DMD, REM

Investigation: NNO, TZ, WB, DMD

Visualization: NNO, TZ, DMD

Funding acquisition: DMD, PM, NNO, TC, TZ

Project administration: DMD, NNO, PM

Supervision: DMD, PM, NNO, TC, TJL, EJAC

Writing – original draft: NNO, DMD

Writing – review & editing: NNO, DMD, EJAC, PM, ERIC, TJL, REM, WB, TC

## Ethical statement

All animal methods detailed in this study were performed under licence from the Home Office (Project Licence 60/4109 to Pat Monaghan) and the Ethical Committee of the University of Glasgow.

## Data, code, and materials availability

The detailed step-by-step protocol to perform the library preparation is publicly available at https://www.protocols.io/view/ovation-rrbs-methyl-seq-library-preparation-protocol-kxygx3kydg8j/v1. The raw sequencing data is deposited at: (link not public yet). The code required to process the methylation data is deposited at https://github.com/NellyOlova/Ovation_RRBS. The code required to build the clocks and perform statistical analyses on the prediction of the clocks are deposited at https://github.com/dmdominoni/zebbieclock. Code used for plotting and analyses of age-correlated CpGs and CORT-DMRs are deposited at https://github.com/NellyOlova/Zebrafinch_Age-CORT.

## Supplementary Materials

### Materials and Methods

#### Sample collection and structure

The samples were collected between 03/10/2011 and 03/10/2022 at the Zoological Research Facility (ZRF), University of Glasgow. For all animals, blood samples were obtained under licence from the UK Home Office by brachial vein puncture and collected in heparinised capillary tubes. They were then centrifuged to separate the red blood cells, which were immediately frozen at −70° C. Three LOTs of zebra finch samples were collected and processed.

##### LOT1

The first lot of blood samples (n = 52, from 17 birds) were obtained from a biobank of samples collected over several years from birds that had been kept throughout their lives in the ZRF in standard conditions, not exposed to any manipulation. The chronological age of the zebra finches spanned from 0 to 3391□days, covering the full expected lifespan of captive zebra finches, and exceeding average lifespan in the wild (*50*). Sexes were balanced across the age range (Fig. 1A). Most birds were sampled longitudinally, seven birds were sampled once, four birds were sampled three times, two birds were sampled four times, three birds were sampled six times, and one bird was sampled seven times.

##### LOT2

We obtained 96 blood samples from 96 healthy zebra finches that were part of a project on circadian rhythms. Briefly, in this experiment birds were assigned to either a control group (n = 47), where they were exposed to regular light-dark cycles (as for all the birds in Run 1), or to an experimental group (n = 49), in which birds were exposed to constant light throughout the 24 hours (light intensity = 500 lux). Each light treatment group included young (age mean + SD: 248 + 11 days for both controls and experimental) and old birds (age mean + SD: 1920 + 158 days for controls and age mean + SD: 1886 + 163 days for experimental). Sexes were balanced across age classes and treatment groups (Fig. 1b). The control group from this study was part of the clock training set, being exposed to the same environmental conditions as the LOT1 birds. The experimental group was used for clock testing, because no effect on epigenetic age nor on longevity was observed from the constant light exposure.

LOT1 + LOT2 controls = AGE cohort, i.e. clock training set (99 samples from a longitudinal set of 64 healthy known-age zebra finches)

##### LOT3 (CORT cohort)

Derived from on ongoing study on parental effects. A group of 48 nestlings were given the same CORT treatment as previously described in the study of the impact on longevity (*24*, *25*). In brief, nests were assigned to a control or experimental group. Nestling corticosterone levels were carefully manipulated to mimic naturally elevated levels due to environmental stressors. In the experimental group (N = 48, average age 29 days, SD = 0.85), levels of circulating corticosterone (CORT, the primary glucocorticoid hormone in birds) were elevated by oral dosage with CORT in a groundnut oil carrier for 17 days, between days 12 and 28 after hatching. On days 12-15, chicks were given 6.2 µg CORT daily (0.124 mg/ml of corticosterone in groundnut oil), administered as two 25 µl doses 5 hours apart. On days 16-28, chicks were given 8.15 µg CORT daily (0.163 mg/ml in groundnut oil), administered as two 25 µl doses 5 hours apart. Control birds were only given the carrier (N = 48, average age 29 days, SD = 0.76). A single blood sample was collected per bird after CORT exposure, in groups spread between days 29 to 32, which provides a cross-sectional 4-day follow up of the post-stress recovery.

The control birds from LOT3 were used for clock testing and selection, while samples from both control and experimental birds were used to measure epigenetic age after early-life developmental stress.

LOT2 treatment + LOT3 controls = Independent clock test set (96 samples from 96 known-age birds).

#### DNA extraction

DNA was extracted from 7 µl of red blood cells using the Puregene Blood Core kit B (Qiagen), following the manufacturer’s protocol, including the optional RNAse I digestion step. DNA concentration and quality were measured using a Nanodrop-8000 spectrophotometer, and further quantified using a Qubit 4 fluorometer prior to library preparation.

#### Library preparation and sequencing

Libraries were prepared with the Ovation RRBS Methyl-seq System (Tecan) following the manufacturer’s instructions, with some modifications. The detailed step-by-step protocol, used for the preparation of all libraries is publicly available (*51*). The ready libraries were checked on an Agilent Tapestation 4200 prior to sequencing, and quantified by fluorometry using the Qubit dsDNA HS assay before library pooling. Fragment size and quantity measurements were used to calculate molarity for each library. Sequencing (100 bp paired or single end) was performed at the Edinburgh Clinical Research Facility Genetics Core on the Illumina NextSeq 2000 platform using the NextSeq 1000/2000 P3 Reagents (300 cycles) v3 Kit and corresponding Primer Kit, including the custom read 1 sequencing MetSeq primer 1 (Tecan). PhiX Control v3 was spiked in each run at a concentration of 10% for LOT1 and 5% for LOTs 2 and 3.

#### Data processing and analysis pipeline

Original fastq files were demultiplexed using Bcl2fastq2 v2.20.0.422 and quality-checked by FastQC v0.12.1 (*52*). Sequencing adapters and low-quality bases were removed with Trimgalore! v0.6.6 (*53*) using Cutadapt v4.6 (*54*) and additional diversity trimming was performed with a custom script provided by Tecan (https://github.com/nugentechnologies/NuMetRRBS). Data purity was assessed with FastQScreen v0.11.4 (*52*). Reads were aligned to the *Taeniopygia guttata* genome (bTaeGut1.4.pri) with Bismark v0.22.3 (*55*) with settings --bowtie2 --sam using bowtie v2.5.3 (*56*) and samtools v1.16.1 (*57*) to an average mapping efficiency of 48% for all three LOTs. Ready sam files were stripped and deduplicated with custom scripts provided by Tecan (https://github.com/tecangenomics/nudup). Methylation calls were extracted with Bismark v0.22.3 and MultiQC (*58*) was used for summarising data from workflow reports. Bisulfite conversion efficiency was 99.43% on average for all three LOTs, as assessed from CH context (where H is A, C, or T) positive calls. The code for the data processing steps, including all custom scripts provided by Tecan, is deposited at https://github.com/NellyOlova/Ovation_RRBS.

#### Methylation quantitation, data partitioning and filtering

We obtained an average of 41 million (M) reads per sample and 63 M CpG calls were extracted per individual on average (Table S1), spanning over ∼8 M covered CpG positions, out of 9.46 M total annotated CpG positions in the zebra finch genome. Bismark output files (cov) were imported in SeqMonk (*43*) over the bTaeGut1.4.pri genome assembly. For the standard single-CpG clock, ‘Read position probe generator’ was used to call each CpG position, for which at least one (1,937,335 M shared CpGs) observation (sequencing calls) was made for every one of the 99 clock training samples, ensuring that the output methylation data will not have missing values (NAs).

For the regional clocks, three approaches were used to generate variable length regions, expanded from Simpson et al 2023 (*27*) (Fig. S2A). 1) Fixed length regions: performed with the ‘Running window probe generator’ to create sliding window tiles with fixed lengths; 2) Fixed CpG regions: defined with the ‘Read position probe generator’ in SeqMonk to create tiles with an equal number of CpG positions in each tile. 3) Cluster-based regions were generated with the R package *dbscan* with different ‘eps’ values (defining the cluster sizes) and subsequently imported in SeqMonk as coordinates (*60*) to generate methylation matrices.

The regional clocks were also filtered based on coverage, testing for variable filtering stringency. All single-site or regional methylation was quantified with the Bisulfite pipeline in SeqMonk, which first averages the methylation calls over each individual CpG position, and after that averages CpG position values within the coordinates of each region. The methylation values generated from all regional and filtering combinations were exported as separate tables via the Annotated probe report in SeqMonk, from which we obtained ‘m x n’ matrices for running the clock algorithms. Finally, for each partitioning approach, we created two matrices: one containing all CpGs/regions, and the other containing a reduced CpG/regions number, where non-age-correlated sites were removed (R^2^ <0.1).

#### Epigenetic clock algorithm

The data were analysed and visualized in R v4.4.1 (R core team, 2024) with the packages *caret 6.0-94* (*61*) and *glmnet 4.1-8* (*62*). We trained an elastic net regression model, i.e. alpha parameter α=0.5. For the lambda value, we created a grid of 80 different lambda values, and let the model find the optimal value. For this we used a leave-one-out cross-validation (LOOCV), with the modification that one individual rather than one sample was left out at each fold, because of repeated sampling of some LOT1 individuals (i.e. number of individuals equals number of validation folds).

We first built models assuming a linear relationship between chronological and epigenetic age. We then built a second set of models, with a logarithmic transformation of the young ages. We followed the approach described by Horvath (*14*, *17*), where the model is trained on transformed values for young ages below a certain age cutoff, and on non-transformed values for adult ages above the age cutoff. The age cutoff is chosen empirically around or above sexual maturity, and we chose 730 days (2 years) as most suitable.

The following function was used, where age was measured in days:

F(age)=log(age+X)-log(730+X) if age <= 730 days;
F(age)=(age-730)/(730+X) if age > 730 days,

where X is a coefficient in days. We tested coefficient values ranging from 1 to 60 days, and a 4-day offset (the “730_4” approach) produced the lowest mean absolute error (MAE) in clocks across all four partitioning methods (Fig. S2B,C). The inverse of the function F is applied to obtain the predicted epigenetic ages as in Horvath 2013, using the predict() function in *caret* (*61*).

Pre-filtering to age-correlated CpGs was also carried out and used in parallel to the non-filtered data. For the log-linear clocks, logarithmic transformation of ages <= 730 days was also applied when computing the Pearson correlation coefficient of each CpG site.

The resulting clock models from each approach were further validated on independent test RRBS data. Regional values for the test data sets were calculated with the BS pipeline in Seqmonk, by averaging the methylation of the CpG positions within each region, without applying any form of coverage and positional pre-filtering prior to applying the model. Test data methylation matrices were prepared from the all-CpG-containing master matrices for each partitioning, after subsetting for each clock’s coordinates. Imputation was applied only where no genuine methylation calls were available for the whole region; individual missing CpG positions within regions were not imputed, only for the Single-CpG matrices, where necessary. Imputed NAs were assigned the methylation mean value of all samples within a matrix per partitioning.

Important note: the clock ‘test set’ was used to apply the final clock models as an independent performance test on previously unseen data. The models were not improved further for each of the 2,146 partitioning input matrices and were final per approach. However, the test set performance results were used in the next step for validation set of the partitioning and workflow parameters, and ultimately, for final selection of the most appropriate model.

Design limitation: the age distribution of the test samples was uneven and heavily weighted toward young individuals, including the controls of the CORT cohort. This generated an inevitable bias in the final clock selection process, which was in part a sought effect considering the imminent clock application on the very young CORT cohort.

#### Statistical analyses on clock performance and age acceleration

We assessed the performance of each clock by measuring the correlation between epigenetic age and chronological age using the Pearson correlation test and calculating R^2^ and MAE.

To assess the impact of CORT treatment on age acceleration of the CORT cohort (LOT3), we used the difference of predicted age versus actual age (delta age) as a response variable in a Gaussian linear mixed effects model. We applied treatment, sex, the interaction between treatment and sex, and mother age as fixed effects, and nest as a random effect to account for family effects, and calculated both marginal (only fixed effects) and conditional (fixed + random effects) R^2^ values (*63*). The datasets and scripts used for this test are available at: https://github.com/dmdominoni/zebbieclock.

#### Genomic features analyses

Genomic features overlap was performed in SeqMonk, and gene annotation was obtained from the official bTaeGut1.4.pri gtf annotation (GCF_003957565.2). Promoters were defined as −1000 bp to +1500 bp of the TSS, exons and introns were obtained from the annotated transcript sequences, and intergenic sequences - with the Interstitial probe generator. Zebra finch CpG island (CGI) annotation was custom generated (link to script upload), with the following criteria, based on the mammalian CGI definition: minimum 55% GC sequence content within a minimum of 200 base pairs (bp) size, and merging CGIs separated by less than 200 bp apart. Feature overlap was defined as ‘overlapping’ for exon, intron and intergenic, and ‘closest’ within 5 kb for transcript (gene) and promoter.

Genome browser views for *CDKN2C*, *ZBTB16* and *LOC115498543* were extracted from SeqMonk as editable svg files.

#### Differential methylation analysis of CORT treatment

For the differential methylation analysis of the early developmental stress experiment (CORT cohort = LOT3 birds), we used 30-CpG genome partitioning windows (probes) with averaged methylation. Methylation was quantified in the same way as for the clock training, with a minimum of 1 call per probe to retain for further analyses, which removed all probes with NAs. DMRs were then identified using logistic regression with multiple testing correction in SeqMonk v1.49.0 and 60,885 regions were identified with p<0.01 and FDR<0.05. From those, only regions with a minimum of 5% averaged methylation difference in either direction (hypo and hyper) across the treatment and control groups were selected for further analyses. This yielded 3,616 DMRs with FDR<0.03, which makes 1.93% of the total number of probes (187,160). Genes were assigned as directly overlapping or closest within 5 kb of the DMR. For the all-DMR heatmap, hierarchical correlation distance clustering was applied to individuals within the control and CORT groups and plotted in RStudio RStudio 2022.02.2 with R v4.3.1.

#### Gene Ontology analyses

GO terms and pathway analyses were performed with Metascape (*64*) using the zebra finch GO database. To overcome the significantly poorer annotation of the zebra finch GO database, for the HPA and eps250_730_4 gene circos overlap, the human GO data base was used with its corresponding orthologs, alongside the zebra finch GO database and both results are shown, either in the main or in supplementary figures. Gene lists included only unique genes with matching human orthologs and excluded predicted avian genes with no known function. Each gene list was supplied together with a custom background gene list, consisting of 12,509 known zebra finch genes with matched human orthologs (Table S3).

#### Age-correlated CpGs: identification and functional annotation

We started with all shared CpG sites of the clock training age-series data (99 samples; 1,937,335 CpGs). Age-correlated CpGs were identified by applying a Pearson correlation between chronological age and individual CpG methylation values, and selecting CpGs with Pearson correlation coefficient |r| ≥ 0.5, which equals R^2^=0.25. This yielded 5,129 age-associated CpGs (Age-CpGs), which is 0.26% of the total CpGs.

Feature overlap was performed in SeqMonk v1.49.0. We mapped each CpG to a gene that was either directly overlapping or the closest within 5 kb. Functional annotation of the known genes was performed in Metascape as described for the CORT-DMRs, with a custom zebra finch background gene list of 12,353 genes and based on the zebra finch GO database.

#### Principal component analysis (PCA) of intersected Age-CORT CpGs

To examine whether CORT-responsive regions exhibit similar methylation changes to those occurring during natural ageing, we intersected the genomic coordinates between the 5129 age-associated CpGs (selected in the AGE cohort, n = 99) and the CORT-responsive DMRs, identified independently in the CORT experiment (CORT cohort, n = 96). This identified an overlap of 115 Age-CpGs located within CORT-DMRs, out of which 111 yielded individual DNAm values in the CORT cohort data and could be used for paired Age-CORT analysis. Methylation at these 111 individual CpGs was used to construct a PCA based exclusively on the AGE-cohort, i.e. with 99 different age points. PC1 captured 48.5% of the variance and showed high correlation with age (r=0.81), whilst PC2 captured 9.6% of variance and showed no correlation with age (r=-0.04). Methylation values of the same 111 Age-CpGs derived from the CORT-treated and control birds were subsequently projected onto this age-derived methylation space. Data scaling was applied. Importantly, inclusion of an individual CpG in this set was determined by its association with age and its genomic location within a CORT-responsive DMR, rather than by evidence that the CpG itself was differentially methylated between CORT and control birds. Each CORT-DMR was defined by 30 individual CpG positions, and there was no prior knowledge if the intersecting Age-CpG was itself differentially methylated and responsive to CORT-treatment.

#### Statistical analyses (non-clock related)

Statistical analyses for the Control vs CORT within-gene comparisons of CORT-DMRs were performed with two-sided t-tests in GraphPad Prism v8 (classical or Welch’s depending on S.D. and distribution).

Observed-versus-expected by chance gene lists overlap was calculated in R v4.3.1 with a one-sided Fisher’s exact test and a Benjamini-Hochberg multiple-comparisons correction to obtain the false discovery rate (FDR).

Pearson correlation on the methylation changes of Age-CORT CpGs across the aging trajectory (YR9 – D1) and CORT response (CORT-Control) was performed in GraphPad Prism v8.

For the PCA analyses, statistical significance between the Control and CORT PC1 groups was assessed with Welch’s t-test (actual p-value = 7.666e-08, Control PC1 mean = 6.707945 and CORT PC1 mean = 9.974523) in R v4.3.1.

Gene methylation changes across the aging trajectory (for *ZBTB16*, *FKBP5*, *NR3C1* and *NR3C2*) were assessed in GraphPad Prism v8 with a one-way ANOVA with Bonferroni multiple testing correction, comparing the means of the CORT data with the Control and all other age groups. Same test was used for the CORT recovery phase (for *ZBTB16*, *FKBP5*, *NR3C1*), where CORT values per recovery day was tested against the pooled Controls. CORT recovery days 3 and 4 (D31 and D32 of sample collection) were pooled for the test to balance replicate number per day.

## Supplementary tables

**Table S1.** Sample metadata and sequencing results (Excel).

**Table S2.** Top Clustering eps250_730_4 clock coordinates and weights (Excel).

**Table S3.** Gene lists used for GO enrichment analyses (Excel).

**Table S4.** Descriptive statistics for the LOT3 CORT cohort predicted age values.

|  | **Control** | **CORT** |
| --- | --- | --- |
| **Number of values** | 48 | 48 |
| **Minimum** | -16.38 | -10.47 |
| **25% Percentile** | -10.04 | -3.688 |
| **Median** | -3.757 | 3.946 |
| **75% Percentile** | 1.813 | 15.18 |
| **Maximum** | 29.45 | 53.42 |
| **Range** | 45.83 | 63.88 |
| **Mean** | -2.625 | 8.490 |
| **Std. Deviation** | 9.604 | 15.86 |
| **Std. Error of Mean** | 1.386 | 2.289 |
| **Lower 95% CI of mean** | -5.414 | 3.884 |
| **Upper 95% CI of mean** | 0.1639 | 13.10 |
| **Coefficient of variation** | 365.9% | 186.8% |
| **Skewness** | 1.395 | 1.089 |
| **Kurtosis** | 2.382 | 0.7246 |

**Table S5.**
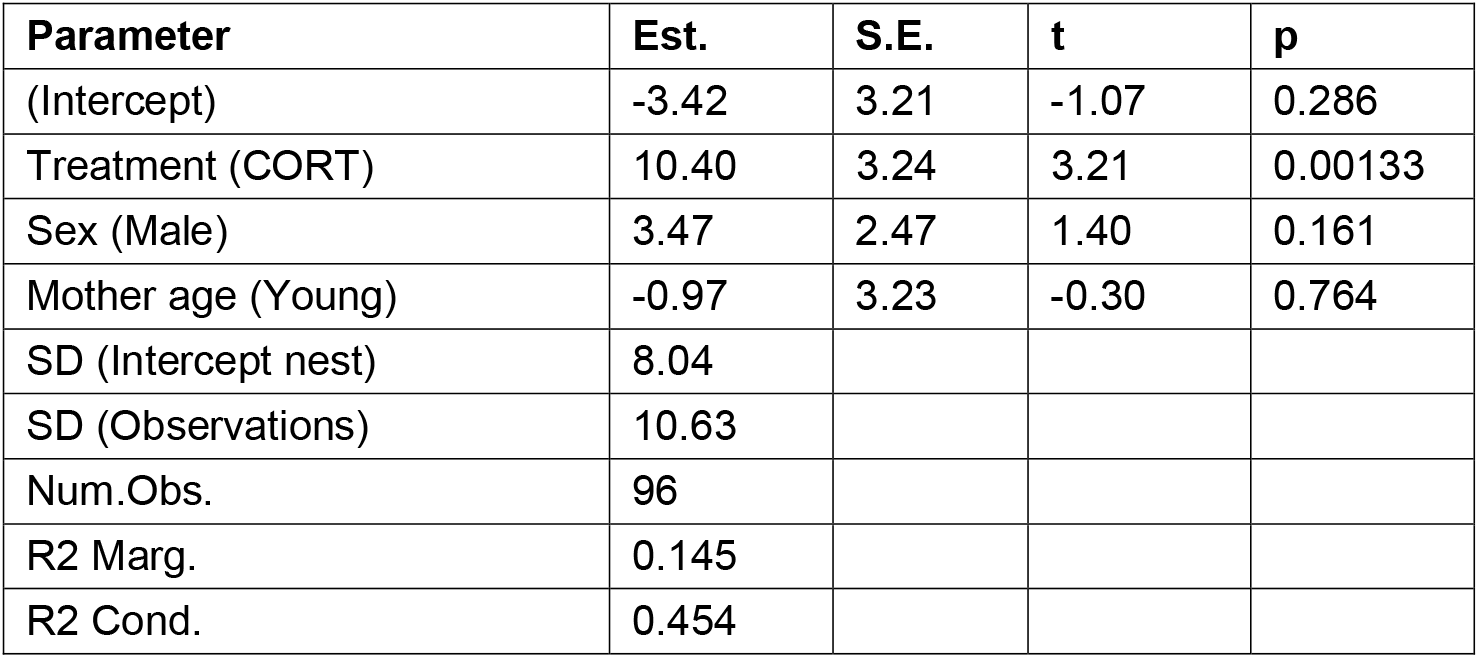
Results of linear mixed models testing for the effects of treatment, sex and mother age on the difference between chronological age and predicted age in the CORT cohort.

**Table S6.** Annotated list of CORT-DMRs (Excel).

**Table S7.** Annotated coordinates of Age-CpGs (Excel).

**Table S8.** Shared Age-CORT CpGs (Excel).

**Figure S1.**
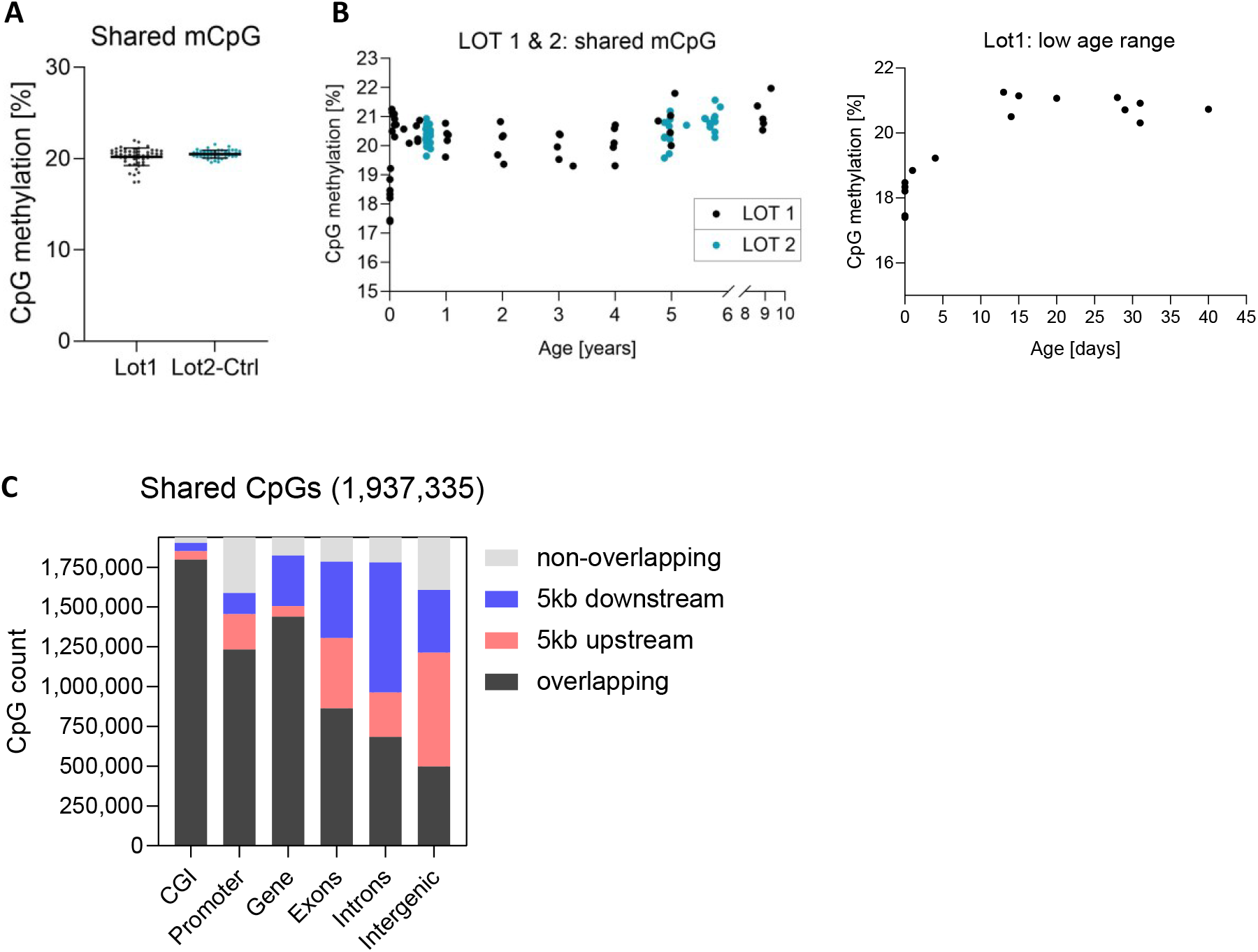
Global DNA methylation and genomic features. (**A**) Global methylation levels of 1,937,335 shared CpG positions across 99 AGE cohort samples used for the clock training; (**B**) Global methylation plotted against the age of individual birds (left); global methylation levels within the first 45 days (right); (**C**) Genomic feature enrichment of the RRBS-derived CpG methylation data.

**Figure S2.**
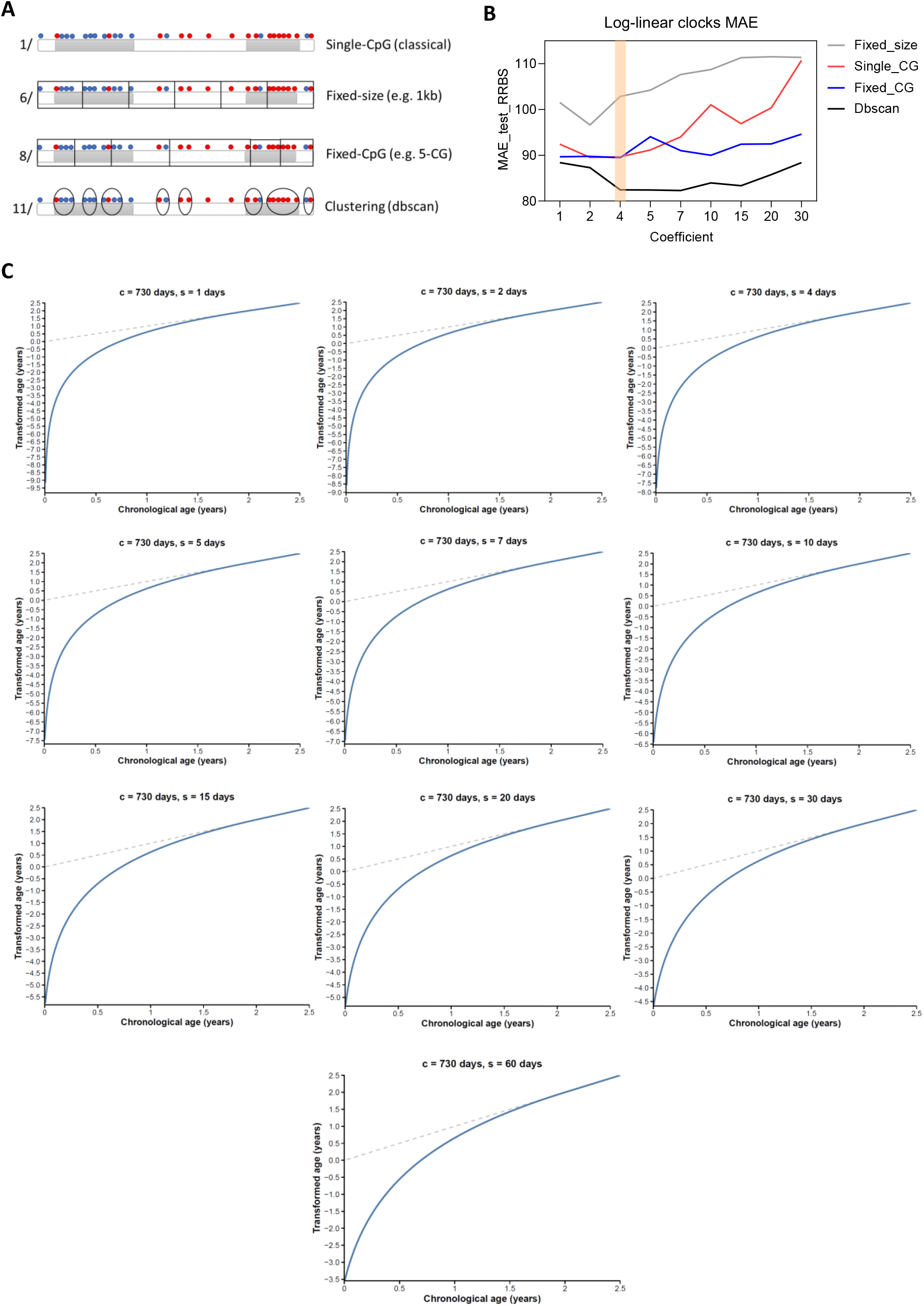
Epigenetic clock construction. (**A**) A schematic of the four genome partitioning methods used for RRBS-based clock construction; (**B**) Tested coefficient values for the logarithmic transformation of ages below 730 days and the transformed ages slopes; (**C**) MAE comparison of the tested coefficient values for the logarithmic transformation of ages below 730 days.

**Fig. S3.**
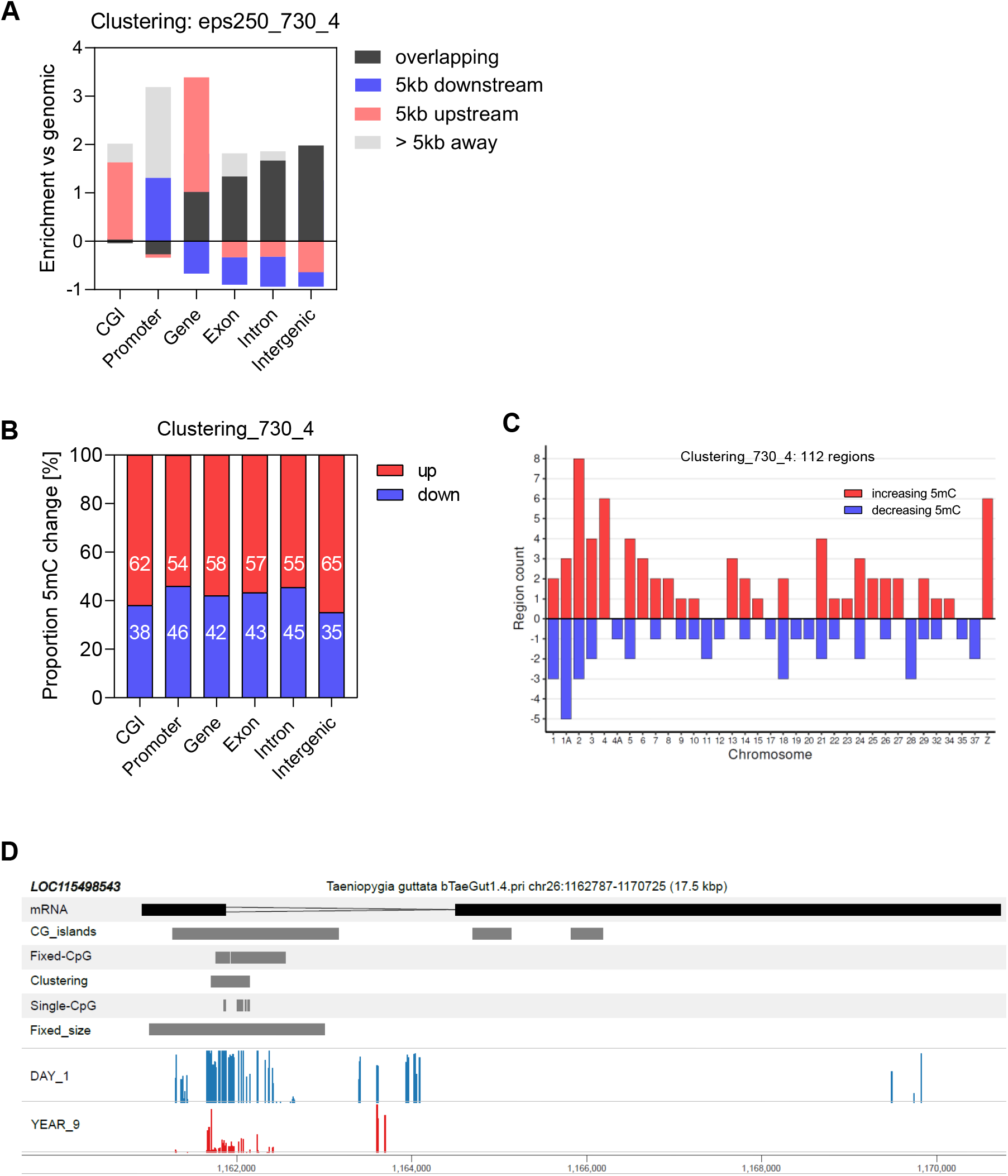
Features of the eps250_730_4 clock. (**A**) Relative feature enrichment over the eps250_730_4 CpG coordinates against the feature distribution over the total 1,937,335 CpGs; (**B**) Proportion of hyper- and hypo-methylated regions within the different genomic features covered by the ‘eps_250_730_4’ coordinates. (**C**) Distribution of the eps250_730_4 clock segments with increasing or decreasing 5mC across the zebra finch chromosomes. (**D**) Chromosome view of the *LOC115498543* ncRNA locus; Clock segments and CGIs are depicted as grey bars, the mRNA is in black, and methylation percentage (5mC %) bars are in blue for ‘day 1’, and in red for ‘year 9’ samples.

**Figure S4.**
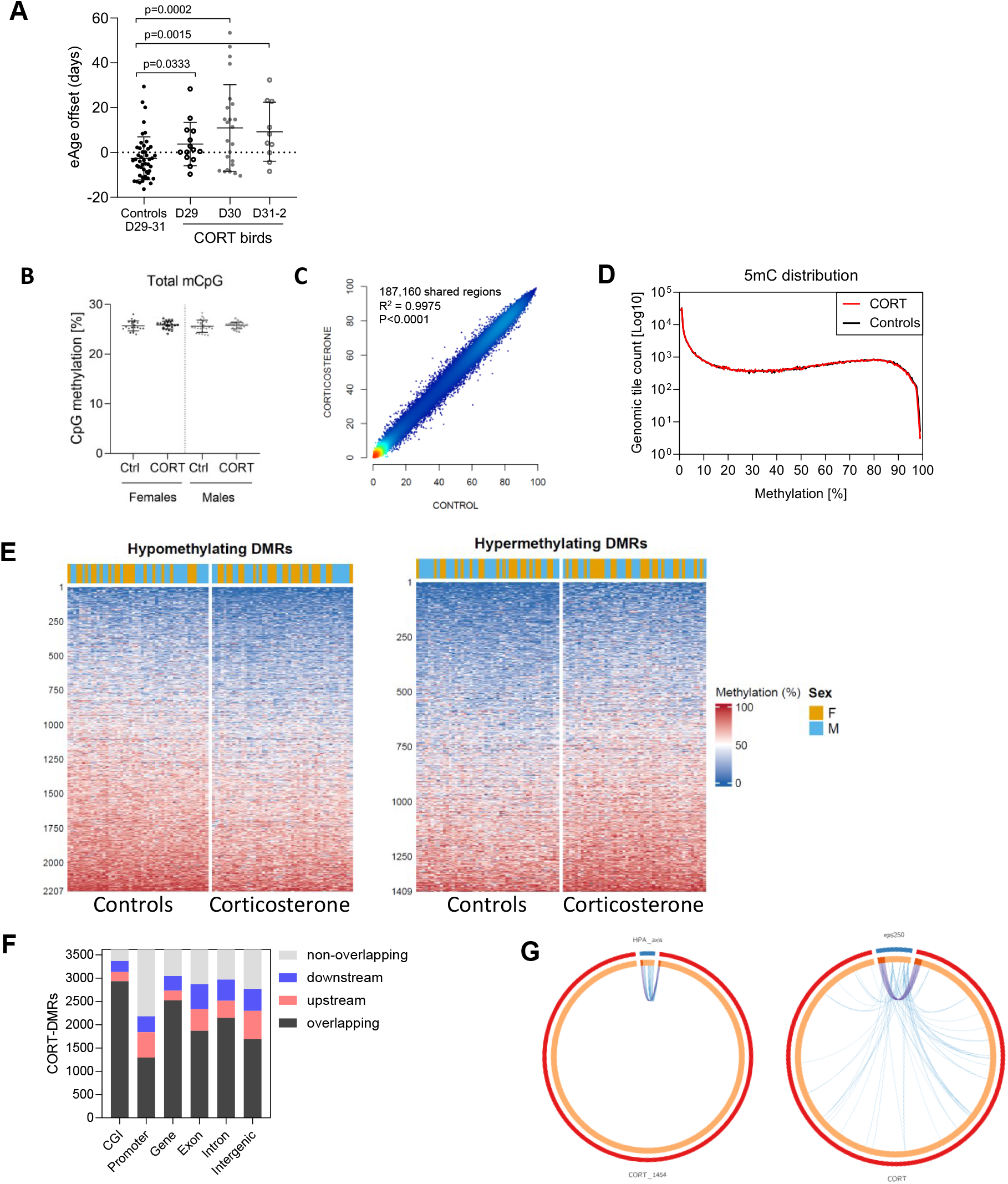
Genomic features of experimental CORT exposure. (**A**) Accelerated epigenetic age across days post CORT treatment; Controls are pooled (days 29-31), CORT results were pooled for days 31 and 32. Significance was assessed with a two-tailed t-test against the Controls, error bars represent S.D. (**B**) Global genomic mCpG levels of CORT-treated samples and controls across males and females. (**C**) Comparison of average 5mCpG methylation across all 187160 shared regions between the CORT and Control groups. (**D**) Distribution of global methylation levels of the CORT and Control groups per genomic tile count. (**E**) Individual CORT-DMR increasing or decreasing methylation levels between CORT and Control samples; the values represent total 5mCpG levels. (**F**) Genomic feature enrichment of the CORT-DMRs by DMR count per each category. (**G**) Circos plots identical to those in Fig.3 but based only on zebra finch gene ontology terms, which are much more limited than the human ontology. Left: depicting direct (shared gene = purple lines) and indirect (shared biological pathway = blue lines) overlap between 1,441 known-function CORT-affected genes and 35 HPA-axis genes. Right: depicting the direct and indirect functional overlap between 1,441 CORT-affected genes and the 84 known-function eps250_730_4 clock-associated genes.

**Figure S5.**
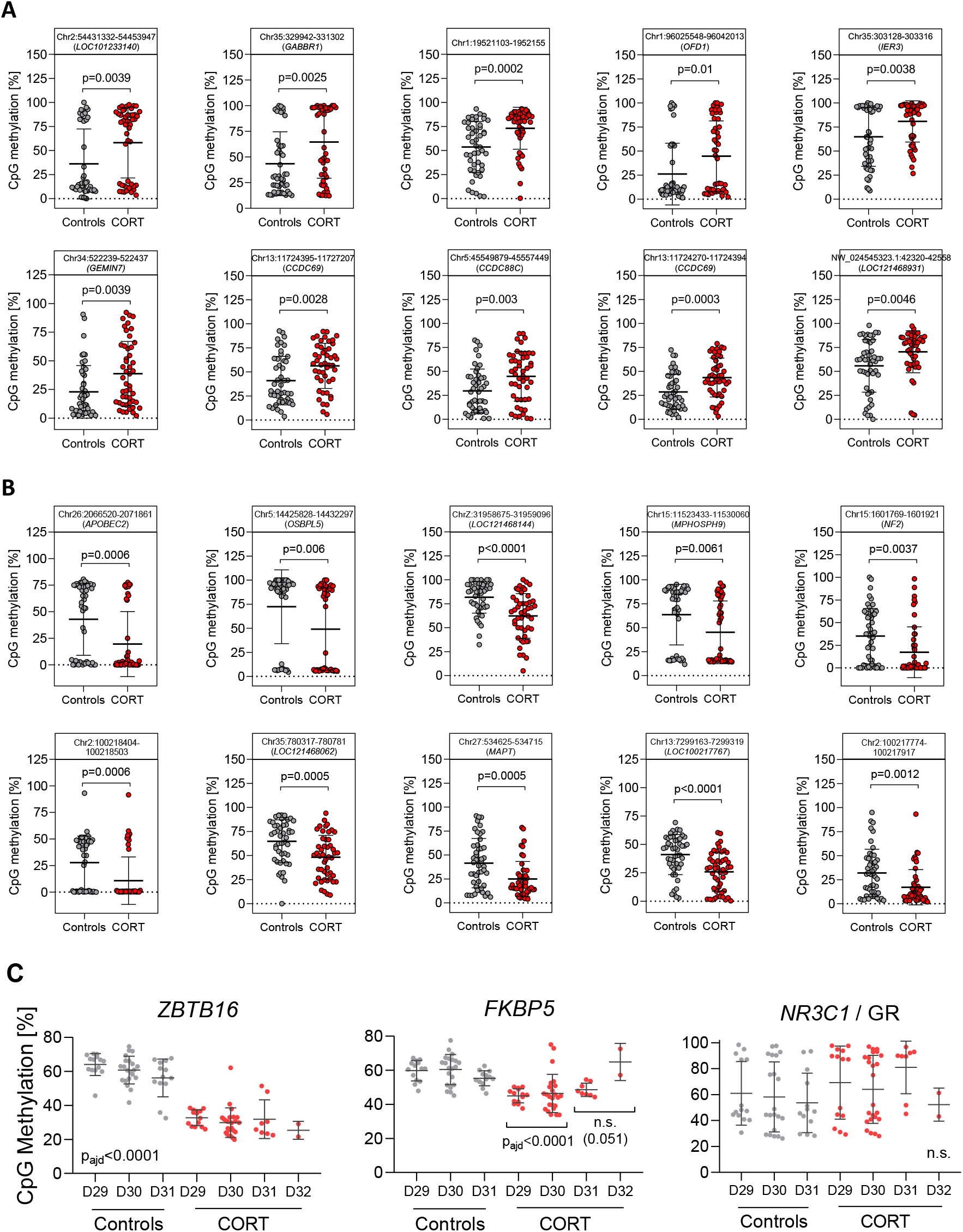
Top hyper- and hypo-methylated CORT-responsive DMRs. (**A**) Top 10 hyper-methylated and (**B**) top 10 hypo-methylated CORT-responsive DMRs, labelled by their coordinates and gene associations, where a gene is located within 5kb of the DMR. *ZBTB16* (Chr24:7290035-7290405), the top hypomethylated DMR, is shown in Fig. 3F and excluded here. Error bars show S.D. over the mean value. Significance is calculated by an unpaired two-tailed t-test. (**C**) CpG methylation dynamics of *ZBTB16*, *FBBP5* and the GR *NR3C1* across controls and treated birds, according to sample collection day after CORT withdrawal (D28). Significance was calculated with a one-way ANOVA, taking all controls as one group irrespective of day, and pooling CORT values for D31 and D32; which are plotted separately for visualitzation.

**Figure S6.**
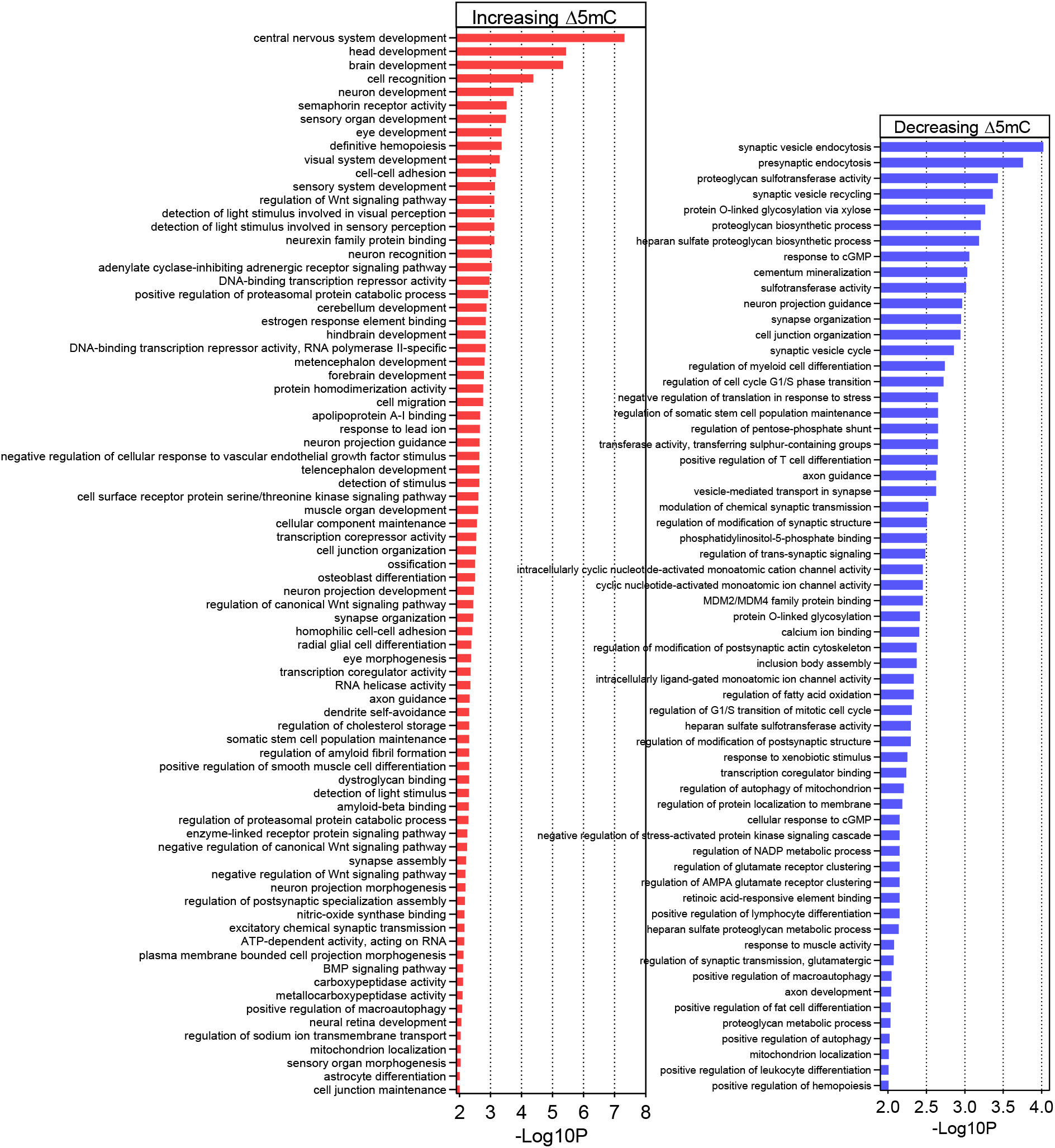
Gene Ontology enrichment for early life CORT exposure. The plots represent all significant (p<0.01) Biological Process terms of hyper-(left) and hypo-methylated (right) CORT-DMR-associated genes.

**Figure S7.**
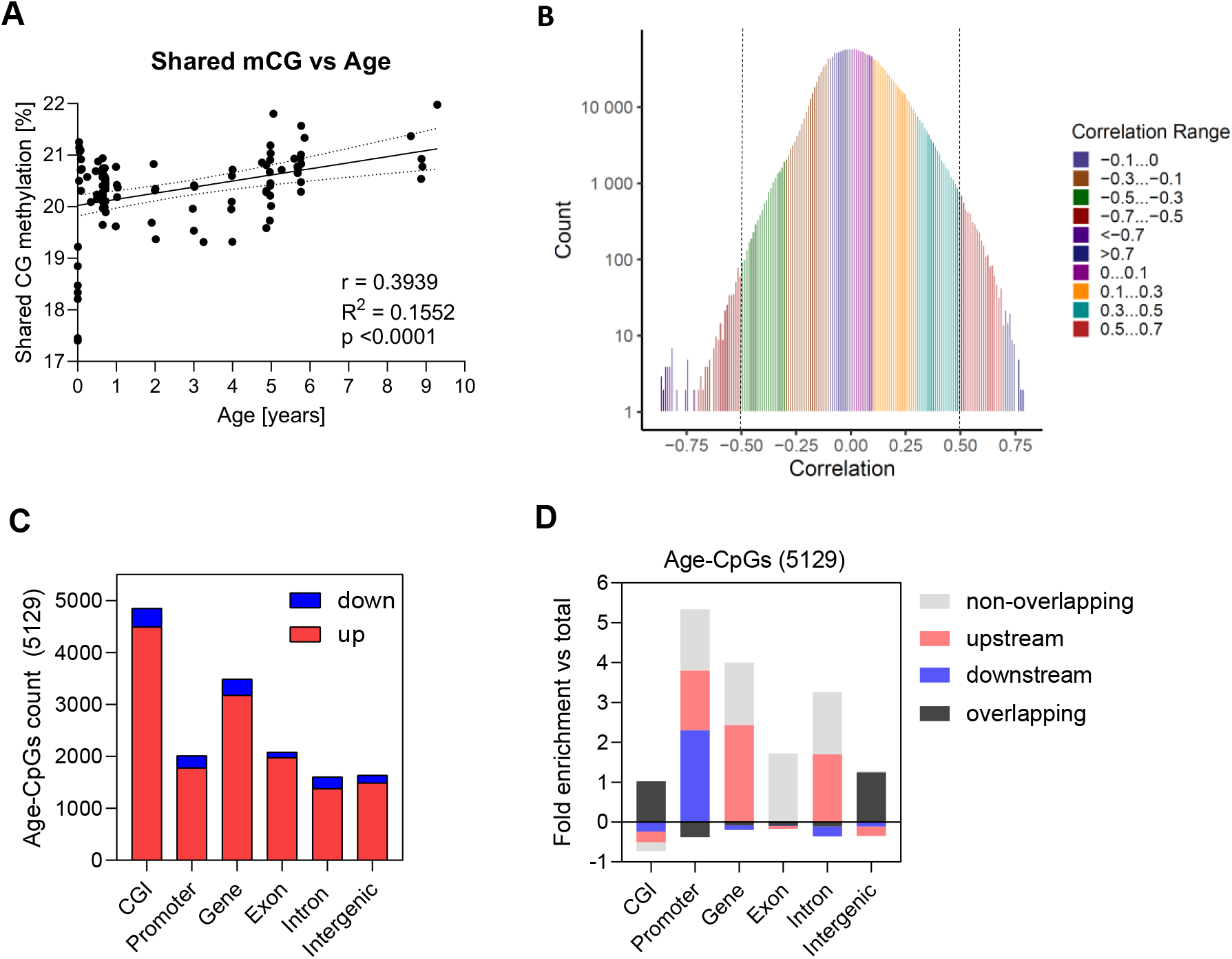
Selection and features of age-correlated CpGs. (**A**) Global genomic 5mCpG levels and their Pearson correlation with age. (**B**) Distribution of the Pearson correlation coefficient r of all 1,937,335 CpGs with the retention criterion cutoff at r=0.5 and r=-0.5; the Y axis is represented as Log(10) of CpG count. (**C**) Proportion of hyper-(up) and hypo-methylated (down) age-associated Age-CpGs per genomic features. (**D**) Relative feature enrichment over the 5,129 Age-CpG coordinates against the feature distribution over the total 1,937,335 CpGs.

**Figure S8.**
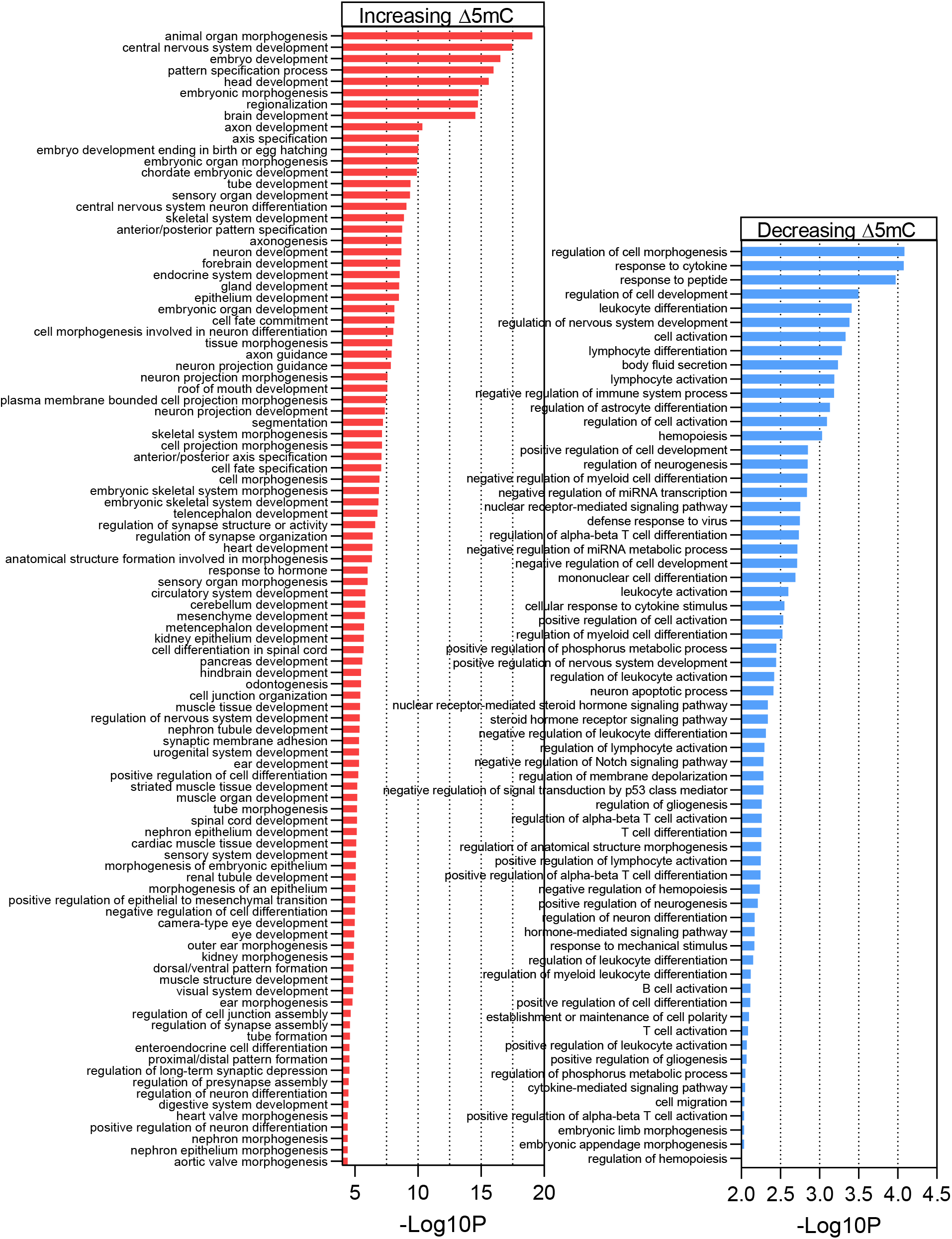
Gene Ontology enrichment for natural aging-associated genes. The plots represent all significant (p<0.01) Biological Process terms of hyper-(left) and

**Figure S9.**
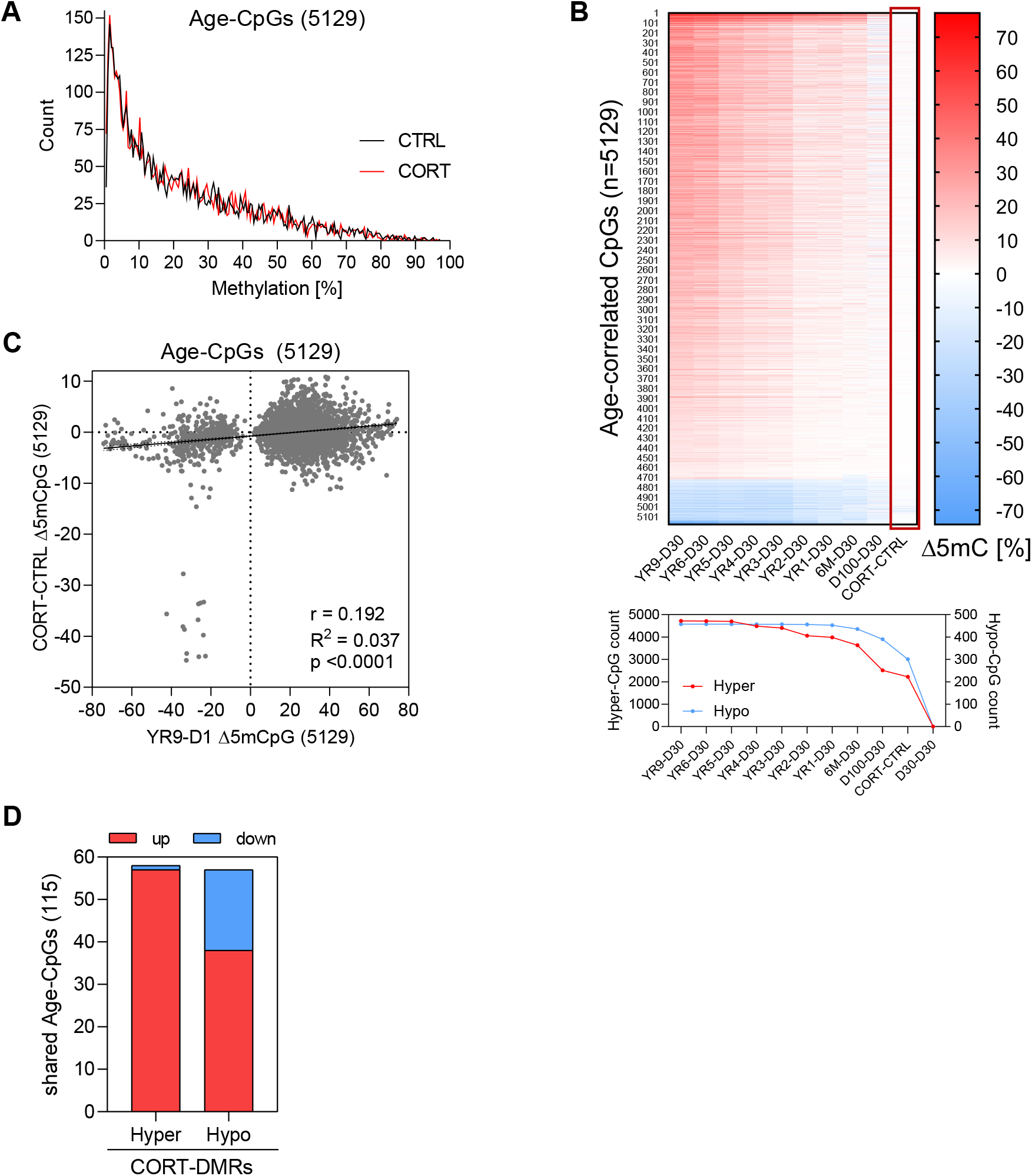
Age-CpGs are affected by early life CORT exposure. (**A**) Distribution of methylation levels of the CORT and Control groups for the 5,129 Age-CpGs. (**B**) Methylation differences between day 30 (D30) samples and all upper age groups alongside the CORT-Control difference (marked in red rectangle, upper plot) and the number of hyper- and hypo-methylated Age-CpGs from D30 onwards alongside the CORT-Control group (lower panel). (**C**) Methylation difference of 5,129 Age-CpGs for the natural aging trajectory (YR9-D1) or upon CORT exposure (CORT-CTRL). (**D**) Direction of the methylation change of 115 Age-CpGs, localised within hyper- or hypo-

**Figure S10.**
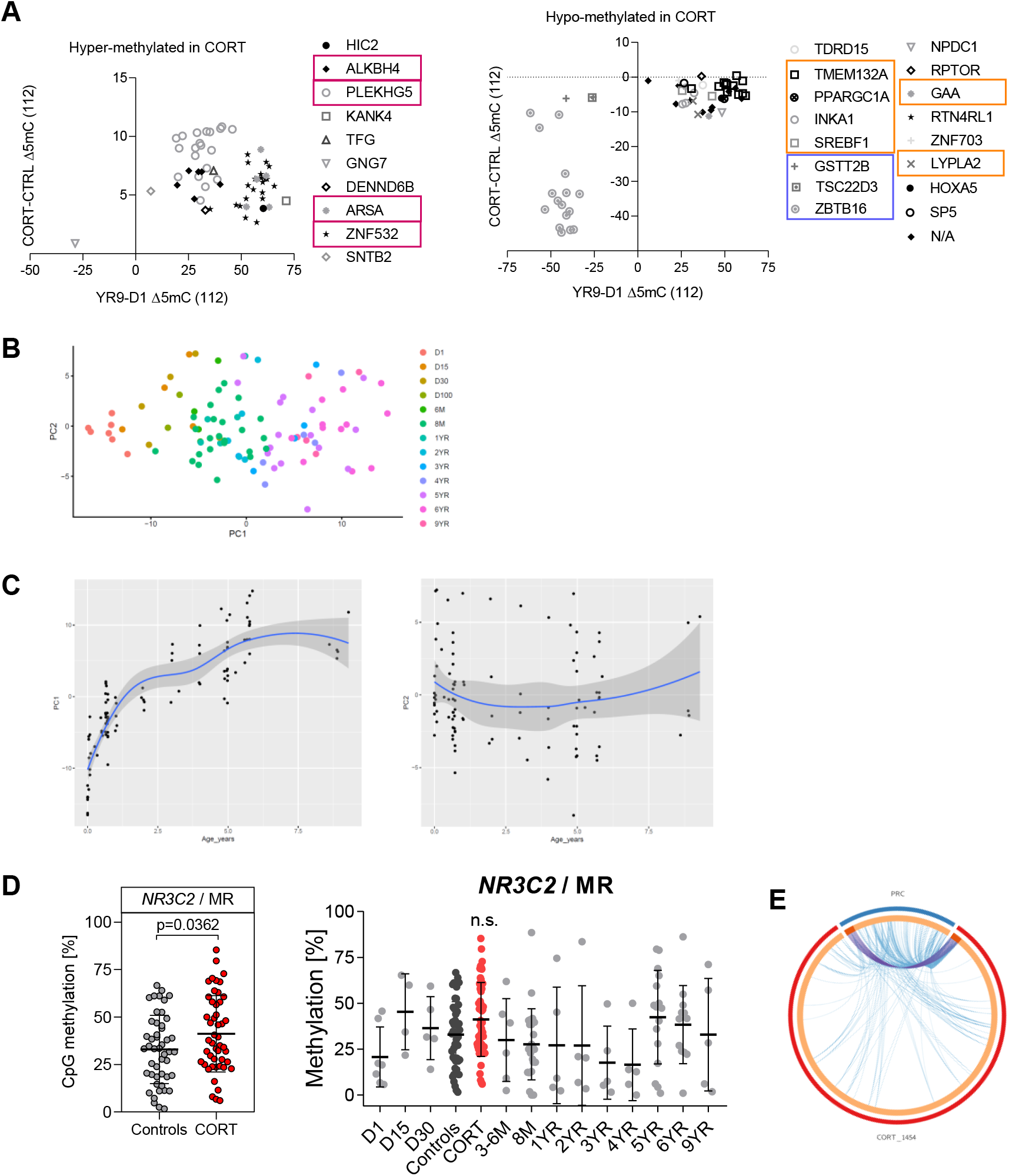
Early life CORT exposure rewires epigenetic pathways related to aging. (**A**) Methylation difference of 115 Age-CpGs overlapping with CORT-DMRs for the natural aging trajectory (YR9-D1) or upon CORT exposure (CORT-CTRL): increasing (left panel) and decreasing (right panel) CORT-DMRs. Each sign represents an Age-CpG annotated or not (N/A) to a gene; genes mapping to multiple CpGs are marked with red or blue rectangles; orange rectangles mark genes with opposite methylation change direction between CORT and aging, relating to enhanced metabolism. (**B**) PCA representing the AGE-cohort (clock-training 99 control samples of different ages) plotted onto the PC space and labelled by age group. (**C**) PC1 (left) and PC2 (right) plotted against chronological age. (**D**) Methylation of the CORT-DMR (Chr4:57888602-57898138) within the mineralocorticoid receptor (MR) *NR3C2* upon CORT exposure (left) and in the context of its normal methylation status within the entire age trajectory (right). (**E**) Direct (shared genes = purple lines) and indirect (shared biological pathway = blue lines) overlap between 1,441 known-function CORT-affected genes and 339 PRC-target genes; Fisher’s p=0.97.

